# Invasion fitness for gene-culture co-evolution in family-structured populations and an application to cumulative culture under vertical transmission

**DOI:** 10.1101/102624

**Authors:** Charles Mullon, Laurent Lehmann

## Abstract

Human evolution depends on the co-evolution between genetically determined behaviors and socially transmitted information. Although vertical transmission of cultural information from parent to offspring is common in hominins, its effects on cumulative cultural evolution are not fully understood. Here, we investigate gene-culture co-evolution in a family-structured population by studying the invasion fitness of a mutant allele that influences a deterministic level of cultural information (e.g., amount of knowledge or skill) to which diploid carriers of the mutant are exposed in subsequent generations. We show that the selection gradient on such a mutant, and the concomitant level of cultural information it generates, can be evaluated analytically under the assumption that the cultural dynamic has a single attractor point, thereby making gene-culture co-evolution in family-structured populations with multigenerational effects mathematically tractable. We apply our result to study how genetically determined phenotypes of individual and social learning co-evolve with the level of adaptive information they generate under vertical transmission. We find that vertical transmission increases adaptive information due to kin selection effects, but when information is transmitted as efficiently between family members as between unrelated individuals, this increase is moderate in diploids. By contrast, we show that the way resource allocation into learning trades off with allocation into reproduction (the “learning-reproduction trade-off”) significantly influences levels of adaptive information. We also show that vertical transmission prevents evolutionary branching and may therefore play a qualitative role in gene-culture co-evolutionary dynamics. More generally, our analysis of selection suggests that vertical transmission can significantly increase levels of adaptive information under the biologically plausible condition that information transmission between relatives is more efficient than between unrelated individuals.

## Introduction

Cultural evolution, which is the change in non-genetically transmitted phenotypes (or information) carried by individuals in a population, is thought to have played a major role in human’s ecological success (e.g., Laland et al., 2010; Boyd et al., 2011; van Schaik, 2016). Cultural evolution rests on mechanisms by which individuals learn and communicate, which themselves depend on behavior or cognitive rules that are at least partially genetically determined. Conversely, cultural evolution can significantly affect reproduction and survival, which in turn affects selection on genes determining behavior. Hominin evolution is therefore influenced by gene-culture co-evolution, whereby genetically determined behavior rules co-evolve along culturally transmitted information (Feldman and Cavalli-Sforza, 1976; Lumsden and Wilson, 1981; Aoki, 1986; Boyd and Richerson, 1985; Feldman and Laland, 1996; van Schaik, 2016).

It is useful to distinguish between two broad cognitive mechanisms that underlie cultural evolution. First, cultural evolution depends on individual learning (IL), which is a generic term for the cognitive processes that lead to the creation of *de novo* non-innate information by an individual, including trial-and-error learning, statistical inference, or insight (Boyd and Richerson, 1985; Rogers, 1988; Dugatkin, 2004; Aoki and Feldman, 2014; Wakano and Miura, 2014). Second, cultural evolution is underlain by social learning (SL), which refers to the cognitive processes that lead to the acquisition of non-innate information from others (Cavalli-Sforza and Feldman, 1981; Boyd and Richerson, 1985; Rogers, 1988; Dugatkin, 2004; Aoki and Feldman, 2014; Wakano and Miura, 2014). If genes are almost always transmitted vertically from parent to offspring, cultural information can be acquired and transmitted in multiple ways via SL. It can be transmitted vertically from parent to offspring, but also horizontally from peer-to-peer, or obliquely between unrelated individuals belonging to different generations (Cavalli-Sforza and Feldman, 1981).

While IL results in the generation of novel information, SL enables the acquisition of skills or information that an individual would be unable to acquire alone by IL over the course of its lifetime. SL thus enables cumulative culture, which is a hallmark of cultural evolution in human populations (e.g., Boyd et al., 2011; van Schaik, 2016). A necessary but not sufficient condition for cumulative culture to occur in a population is that individuals use a composite learning strategy in which SL precedes IL (Boyd and Richerson, 1985; Enquist et al., 2007; Aoki et al., 2012).

Since both IL and SL strategies determine cultural evolution, much theoretical population work on gene-culture co-evolution has been devoted to understand the co-evolution between genetically determined IL and SL learning on one hand, and the amount of cultural information they generate in evolutionary stable population states on the other. This has led to a rich literature investigating the role of various factors, such as the type of cultural information, the regime of environmental change, or the structure of the population for the evolution of IL and SL and their impact on cumulative culture (e.g., Boyd and Richerson, 1985; Feldman et al., 1996; Wakano et al., 2004; Wakano and Aoki, 2006; Enquist et al., 2007; Rendell et al., 2010; Nakahashi, 2010; Aoki et al., 2012; Lehmann et al., 2013; Nakahashi, 2013; Aoki and Feldman, 2014; Wakano and Miura, 2014; Kobayashi et al., 2015; Ohtsuki et al., 2017).

The vast majority of such theoretical work has focused on horizontal and oblique transmission, assuming that information is transmitted independently from genotype via SL (i.e., between randomly sampled individuals in the population, from the same generation for horizontal transmission and different generations for oblique transmission). In this case, the fate of a mutant strategy, here a genetically determined phenotype affecting IL and SL, that arises as a single copy in a resident population can be studied under the assumption that the mutant is in a cultural environment determined only by the resident (i.e., “evolutionary invasion analysis”). In other words, SL among mutants is so rare that it can be neglected (no mutant-mutant interactions), which greatly simplifies mathematical analysis (see Metz, 2011 for general consideration on evolutionary invasion analysis and Aoki et al., 2012 for applications to cultural evolution).

When information is transmitted vertically via SL, however, non-random interactions occur between individuals as transmission occurs in a way that is correlated to genotype. As a result, transmission of cultural information among mutants can no longer be neglected and influences the fate of a mutant strategy. In other words, kin selection occurs (i.e., natural selection when individuals interact with others that are more likely to share a recent common ancestor than individuals sampled randomly from the population, Hamilton, 1964; Michod, 1982; Frank, 1998; Rousset, 2004; van Baalen, 2013; Lehmann et al., 2016). Intuitively, kin selection will favor learning strategies that promote the transmission of adaptive information since related individuals in downstream generations preferentially benefit from this information. This should influence cultural transmission from parent to offspring and hence the selection pressure on IL and SL, which will depend on multigenerational effects of genetically determined phenotypes.

Despite the potential importance of vertical transmission and its prominent role in cultural evolution theory (e.g., Cavalli-Sforza and Feldman, 1981; Feldman and Zhivotovsky, 1992; McElreath and Strimling, 2008), few studies have investigated the evolution of IL and SL and its effect on cumulative culture under vertical transmission. Models with vertical cultural transmission either do not allow for the transformation of cultural information by genetically determined phenotypes (and therefore do not allow to consider cumulative cultural evolution with multigenerational effects, e.g., Feldman and Zhivotovsky, 1992); or make specific assumptions on learning dynamics and the trade-off between resource allocation into learning and into reproduction (Kobayashi et al., 2015; Ohtsuki et al., 2017), from which it is difficult to get a broad view of the impact of vertical transmission on cultural evolution. The learning-reproduction trade-off, which reflects biologically realistic constraints between life-history components, can significantly compromise the accumulation of culture (Nakahashi, 2010; Lehmann et al., 2013; Wakano and Miura, 2014; Kobayashi et al., 2015; Ohtsuki et al., 2017). However, its general importance is not fully understood because most previous studies assume that the marginal cost of learning (i.e., the effect of learning on reproductive success holding everything else is constant), which influences the learning-reproduction trade-off, is constant (Lehmann et al., 2013; Wakano and Miura, 2014; Kobayashi et al., 2015; Ohtsuki et al., 2017, but see Nakahashi, 2010 for an exception).

The above considerations highlight that currently, there exists no framework to systematically carry out an evolutionary invasion analysis for gene-culture co-evolution under vertical transmission with multi-generational effects (i.e., gene-culture co-evolution family structured populations). Since a primary form of transmission of information in humans is vertical (Cavalli-Sforza and Feldman, 1981; Guglielmino et al., 1995; Hewlett et al., 2011; Konner, 2010), such a framework would be useful to understand cultural evolution. In particular, it would allow determining how the level of culture generated by evolving IL and SL departs under vertical transmission from oblique and horizontal transmission, and assess the importance of the learning-reproduction trade-off.

The aim of this paper is therefore two-fold: (1) develop a mathematical model to perform evolutionary invasion analysis for gene-culture co-evolution in a diploid family-structured populations; and (2) study the effects of vertical transmission on the evolution of IL and SL and the concomitant level of adaptive information they generate. In the first part of this paper, we derive the invasion fitness of alleles in a diploid, family-structured populations, when each individual carry cultural information, which can be a stock of knowledge, skill, or any other form of biotic or abiotic capital. The cultural information of an offspring depends deterministically on the cultural information in the parental generation, and therefore on transmission modes, but also on the alleles in the offspring and in the parental generation. In turn, cultural information in the population affect the reproductive success of individuals, resulting in eco-evolutionary feedbacks between genes and cultural information. Second, we apply our framework to the evolution of IL and SL and the concomitant level of adaptive information they generate.

## 1 Gene-culture co-evolution in a family-structured population

### 1.1 Life-cycle

We consider a diploid monoecious population of large and constant size (large enough to neglect random genetic drift) that is structured into families, each founded by a mated individual. The discrete time life-cycle of this population is as follows. (1) Each individual produces offspring and then either survives or dies (independently of age so that there is no explicit age-structure). (2) Random mating among juveniles occurs and then each juvenile either survives or dies (possibly according to density-dependent competition) to make it to the next generation of adults.

Each individual carries a cultural information variable *ε*, which is possibly multidimensional and which can represent the total amount of knowledge or skill held by that individual at the stage of reproduction. The cultural information variable *ε* is indirectly influenced by the genetic composition of the population and affects the survival of adults or juveniles, and/or individual fecundity.

### 1.2 Evolutionary invasion analysis

We assume that only two alleles can segregate in the population, a mutant with type *τ* and a resident with type *θ*, which are taken from the set Θ of all possible types. In order to determine the fate (establishment or extinction) of the mutant *τ* when it arises as a single copy in the population (e.g., Fisher, 1930; Hamilton, 1967; Maynard Smith, 1982; Eshel and Feldman, 1984; Charlesworth, 1994; Metz et al., 1996; Ferrière and Gatto, 1995; Metz, 2011; van Baalen, 2013), we seek an expression for invasion fitness *W* (*τ, θ*), which is here taken to be the *geometric growth rate* of the mutant *τ* introduced into the *θ* population (Cohen, 1979; Tuljapurkar, 1989; Caswell, 2000; Tuljapurkar et al., 2003). Invasion fitness *W* (*τ, θ*) is the per capita number of mutant copies produced asymptotically over a time step of the reproductive process by the whole mutant lineage descending from the initial mutant (as long as the mutant remains rare), and if *W* (*τ, θ*) is less or equal to one, then the mutant lineage goes extinct with certainty (Lehmann et al., 2016). Therefore an uninvadable strategy *θ*; namely, a strategy resisting invasion by any type satisfies

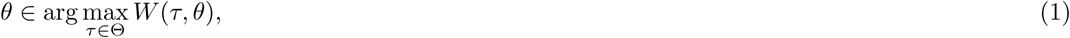

i.e., *θ* maximizes invasion fitness for a resident population at the uninvadable state.

In order to understand gene-culture co-evolution, we aim to derive an expression for invasion fitness *W*(*τ, θ*) that incorporates how gene (embodied by the segregating alleles, which for example code for genetically determined learning strategies) and culture (embodied by cultural information *ɛ*) both affect the reproductive success of individuals, and how *ɛ* depends on alleles in the population. In this task, we first note that under the assumption of random mating, we can neglect homozygote mutants since they are rare during invasion and so only need to consider mutant individuals that are heterozygotes. Then, we denote by *w*(*τ, θ, ε_t_*(*τ, θ*)) the expected number of successful offspring produced by a heterozygote mutant adult over one life cycle iteration at demographic time *t* = 0, 1, 2, …, since the appearance of the mutant at *t* = 0. Thus, this fitness is an expression for *individual fitness* as defined in population genetics and social evolution (e.g., Nagylaki, 1992; Frank, 1998; Rousset, 2004; Lehmann et al., 2016). Individual fitness *w*(*τ, θ, ε_t_*(*τ, θ*)) depends on both the mutant *τ* and resident *θ* types since (i) heterozygotes carry both alleles and (ii) fitness depends on the behavior of individuals in the population at large, composed of homozygotes for the resident. Individual fitness *w*(*τ, θ, ε_t_*(*τ, θ*)) also depends on the cultural variable *ε_t_*(*τ, θ*) of an individual carrying the mutant at demographic time *t*, which itself depends on *τ* and *θ*.

Since individuals carrying the mutant allele are all heterozygotes during invasion, there are no fluctuations of allele frequencies within mutant individuals (no stochastic fluctuations between heterozygote and homozygote states owing to segregation). Hence, a heterozygote parent can at most have a heterozygote offspring for the mutant allele, and assuming that there are no exogenous stochastic effects on *ε*, we can let the sequence of environments 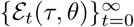 of a lineage of heterozygote mutants follow a discrete deterministic dynamic:

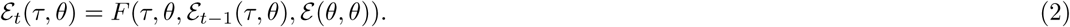

Here, *F* is the cultural map: it transforms the cultural information *ε_t−1_* (*τ, θ*) of a mutant lineage member at generation *t* − 1 into a new cultural information experienced by a lineage member at the next generation *t*. As shown by eq. (2), the cultural information of a mutant depends on its genetically determined behavior (first argument in *F*, *τ*) and that of a resident (second argument in *F*, θ), as well as on the cultural information *ε_t−1_* (*τ, θ*) of a mutant in the parental generation since vertical transmission can occur (third argument in *F*, *ε_t−1_* (*τ, θ*)) and on the cultural information in the resident population (fourth argument in *F*, *ε*(*θ, θ*)) that satisfies

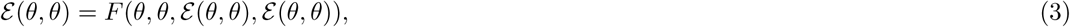

the steady-state version of eq. (2) with *τ* = *θ*. A concrete example for the cultural map *F* within the context of social and individual learning can be found in section 2 (eq. 24).

We assume that the sequence of cultural information 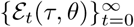 converges to a unique fixed point *ε*(*τ, θ*) = lim_t→∞_ *ε_t_* (*τ, θ*), satisfying

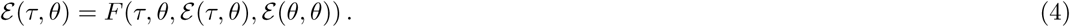

This is the key simplifying assumption in our analysis, from which it follows (see Appendix A) that invasion fitness is equal to

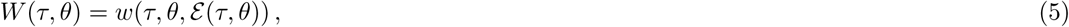

i.e., the individual fitness of a mutant evaluated at the cultural equilibrium of the mutant lineage cultural dynamics (eq. 4).

Invasion fitness *W*(*τ, θ*) as given by eq. (5) shows that it depends on the equilibrium cultural state *ε*(*τ, θ*), which is itself a function of the mutant strategy. Hence, even though the mutant is globally rare, individuals carrying the mutant allele affect the cultural information to which other mutant carriers are exposed in subsequent generations. This results in carry-over effects across generations that influence selection on the mutant.

### 1.3 Selection gradient and local uninvadability

In order to better understand how carry-over effects across generations influence selection, we assume that the genes carried by an individual determine a vector of *n* quantitative traits (Θ = ℝ^*n*^) and that genes have additive effects on phenotype. We let the vector *θ* = (*θ*_1_, *θ*_2_, …, *θ_n_*) ∊ ℝ^*n*^ represent the genetically determined phenotype of a resident individual (homozygote for the resident allele). Similarly, *τ* = (*τ*_1_*, τ*_2_, …, *τ_n_*) ∊ ℝ^*n*^ is the phenotype of a heterozygote individual, and *z* = (*z*_1_*, z*_2_, …, *z_n_*) ∊ ℝ^*n*^ the phenotype of a homozygote mutant. Owing to additive gene action, the phenotypic trait *i* of a heterozygote is the mid-value

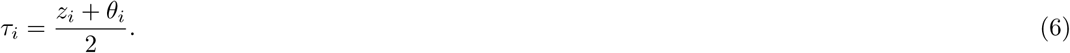

We can therefore write the phenotypic vector *τ* of a heterozygote as *τ* (*z, θ*), which emphasizes that it depends on the two homozygote phenotypes. With this notation, invasion fitness (eq. 5) can be written as

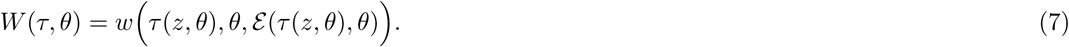

It follows from eq. (7) and the linear dependence of *τ* on *z* and (eq. 6) that

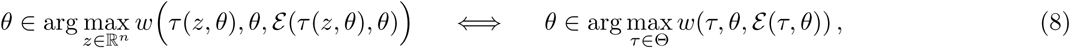

i.e., *θ* is an uninvadable strategy if we cannot find a genetically determined phenotype *z* expressed by a homozygote individual that would make a heterozygote individual with phenotype *τ* (*z, θ*) better off than the resident homozygote *θ*.

The first-order necessary condition for uninvadability must therefore satisfy

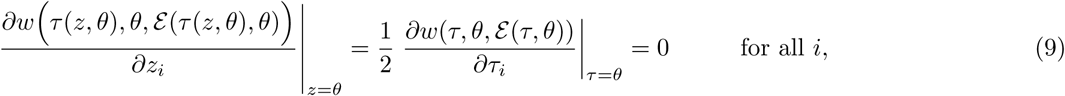

using the chain rule and eq. (6). Hence, we can write the first order necessary condition for uninvadability as

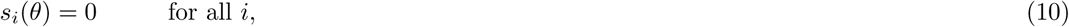

where

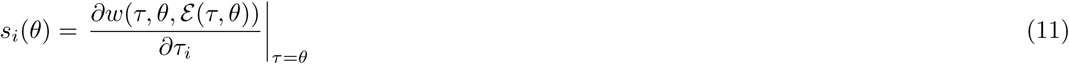

is the selection gradient on trait *i*, which captures the effect of directional selection on trait *i* and allows to compute candidate uninvadable strategies, the so-called singular strategies that satisfy eq. (10) (e.g., Geritz et al., 1998; Rousset, 2004; Leimar, 2009). Eq. (11) shows that under additive gene action (eq. 6), the selection gradient can be expressed in terms of variation in heterozygote effects (the *τ_i_*’s), which has the attractive feature of allowing us to keep simple notations throughout and apply the model to both haploids and diploids with the same notation.

We now disentangle the role of the effects of an individual on its own cultural information (direct effects) from those it has on downstream generations (indirect effects) for selection. To do so, we first use the chain rule to express the selection gradient (eq. 11) as

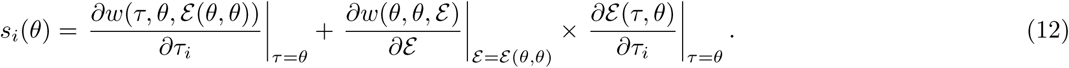

The last term of eq. (12) can be found by first differentiating eq. (4) on both sides with respect to *τ_i_* using the chain rule,

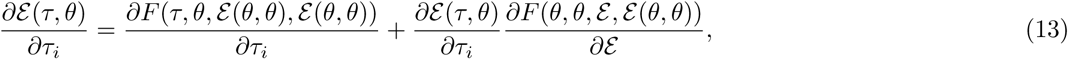

where the derivative in *∂F* (*θ, θ, ε, ε* (*θ, θ*))*/∂ε* is with respect to the third argument of *F* (eq. 4). We then solve eq. (13) for *∂ε* (*τ, θ*)/*∂τ_i_*, which yields

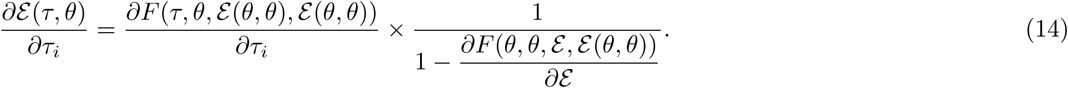

Substituting eq. (14) into eq. (12) gives

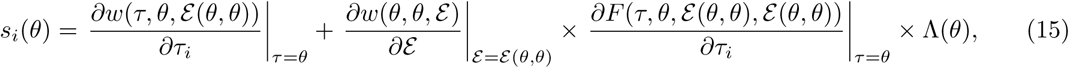

where

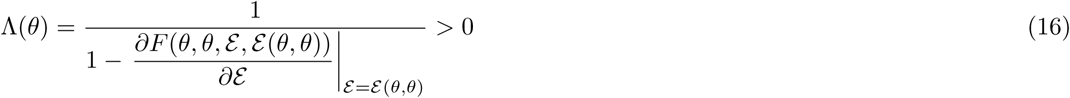

is greater than zero owing to our assumption that the cultural dynamic eq. (2) has a single fixed point (so that −1 *< ∂F* (*θ, θ, ε, ε* (*θ, θ*))*/∂ε <* 1).

Eq. (15) shows that the selection gradient is the sum of two terms. The first terms is the change in the fitness of an individual changing its trait *i* by an infinitesimal amount (i.e., by switching from the resident to mutant trait expression), which is the standard selection gradient in panmictic populations without class structure or effects on cultural information (e.g., Geritz et al., 1998; Rousset, 2004). The second term in eq. (15) captures the fitness effects of cultural changes, cumulated over the lineage of a focal mutant. These cumulative effects are equal to the product of (i) the sensitivity of fitness to cultural change (*∂w*(*θ, θ, ε*)*/∂ε*), (ii) the sensitivity of current cultural dynamics to a change in trait value in an individual (*∂F* (*τ, θ, ε*(*θ, θ*)*, ε*(*θ, θ*))/*∂τ_i_*), and (iii) the multi-generational effects Λ(*θ*) on the culture experienced by a focal mutant of a change in cultural dynamics over all individuals in a line of descent connected to the focal mutant.

The multi-generational effects Λ(*θ*) can be decomposed between the “direct effect” of a focal mutant on its own culture, and the “indirect carry-over effects” of a lineage of mutants on the culture experienced by its members as follows

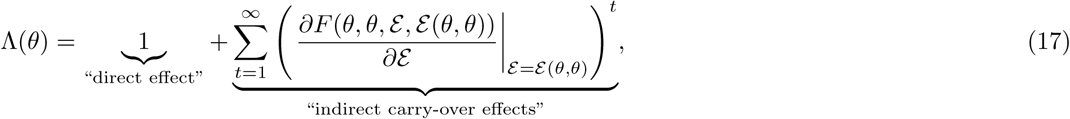

where (*∂F/∂ε*)^*t*^ can be interpreted as the effect of the focal mutant on the culture of an individual living *t* ≥ 1 generations downstream. Hence, in the absence of carry-over effects across generations (*∂F/∂ε* = 0), the selection pressure is given by eq. (15) with Λ(*θ*) = 1. Otherwise, the selection pressure on a trait will depend on its carry-over effects across generations within the family. The selection gradient (eq. 15) can be understood as an inclusive fitness effect of expressing the mutant allele (Hamilton, 1964; Frank, 1998; Rousset, 2004), in which the second summand in eq. (17) is conceptually analogous to the carry-over effects that arise in spatially structured populations when individuals affect local environmental dynamics in a deterministic way (Lehmann, 2008, e.g., eq. 17).

When the difference between non-singular resident and mutant phenotypes is small (∥*τ − θ*∥ ≪ 1), the selection gradient is sufficient to determine whether the mutant will go extinct or fix in the population (see Rousset, 2004 for a general argument about this). A singular phenotype *θ\** (such that *s_i_*(*θ\**) = 0 for all *i*) will then be approached by gradual evolution irrespective of the effects of local mutations on traits, i.e., is strongly convergence stable (Leimar, 2009), if the *n × n* Jacobian J(*θ\**) matrix with (*i, j*) entry

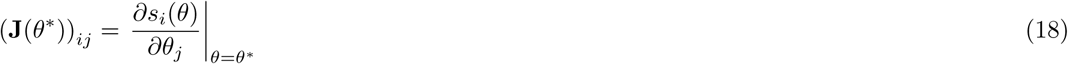

has eigenvalues with negative real parts (or equivalently, if the matrix (J(*θ\**) + J(*θ\**)^T^)/2 is negative-definite, which is the condition given in Leimar, 2009). At a convergence stable singular resident (*s_i_*(*θ\**) = 0 for all *i*), the *n × n* Hessian H(*θ\**) matrix with (*i, j*) entry

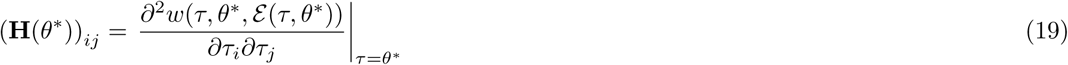

determines whether the resident is locally uninvadable, which is the case if H(*θ\**) is negative-definite at *θ\**. If that is not the case, then disruptive selection and evolutionary branching can occur. Eqs. (18)–(19) give the standard multidimensional (strong) convergence stability and local uninvadability conditions for a finite number of quantitative traits (e.g., Lessard, 1990; Leimar, 2009; Mullon et al., 2016). Further, the eigenvectors of the Hessian matrix can be used to determine which type of correlation between traits will be favored by selection when it is disruptive (Mullon et al., 2016, eq. 9).

## 2 Gene-culture co-evolution and IL and SL levels

### 2.1 Reproductive assumptions

We now apply our framework to study gene-culture co-evolution through IL and SL. We let *ε ∊* ℝ_+_ stand for the amount of adaptive non-innate information that an individual acquires during its lifespan by IL and SL. We are interested in the evolution of resource allocation to IL and SL, and assume that a homozygote individual expressing the resident allele allocates a fraction *θ*_L_ ∊ [0, 1] of its resources to learning (baseline unit of one), and a fraction *θ*_IL_ ∊ [0, 1] of that effort to IL. Hence, an individual allocates *θ*_L_*θ*_IL_ unit of resources to IL, *θ*_L_(1 − *θ*_IL_) units to SL, and 1 − *θ*_L_ to any other function of the organism (e.g., offspring production, maintenance, etc.), creating a trade-off between allocating resources to learning and other functions of the organism (e.g., Nakahashi, 2010; Lehmann et al., 2013; Wakano and Miura, 2014; Kobayashi et al., 2015). With this, a resident homozygote expresses the vector *θ* = (*θ*_IL_, *θ*_L_) ∊ [0, 1]^2^ of phenotypes and a heterozygote mutant has trait vector *τ* = (*τ*_IL_*, τ*_L_) ∊ [0, 1]^2^.

We aim to assess the role of vertical transmission on the uninvadable learning strategy and concomitant level of adaptive information. In order to do this for a mathematically tractable model, we assume that after IL and SL have been performed, an individual gathers energy according to the amount of adaptive information it has learnt and the resources it has left, reproduces using its gathered energy, and then dies (semelparous reproduction). Under these assumptions, the fitness of a heterozygote mutant can be written as

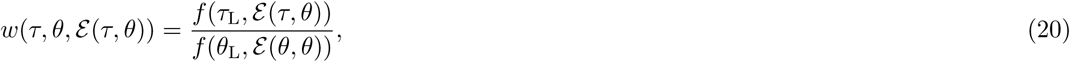

where *f*(*τ*_L_*, ε*(*τ, θ*)) is the fecundity (number of offspring produced) of a mutant, and *f* (*θ*_L_*, ε*(*θ, θ*)) is the average fecundity in the population, which is given by the fecundity of a resident in a monomorphic resident population. Eq. (20) immediately shows that in a resident monomorphic population, invasion fitness is equal to one: *w*(*θ, θ, ε* (*θ, θ*)) = 1.

### 2.2 Learning-reproduction trade-off and learning costs

Since allocating more resources *τ*_L_ into learning deviates resources from reproduction, we assume that fecundity is monotonically decreasing with *τ*_L_ (i.e., *−∂f*(*τ*_L_*, ε*(*θ, θ*))*/∂τ*_L_ *>* 0, which is the “marginal cost of learning”). The trade-off between allocating resources to learning, which is captured by the total derivative of fecundity with respect to learning *τ*_L_,

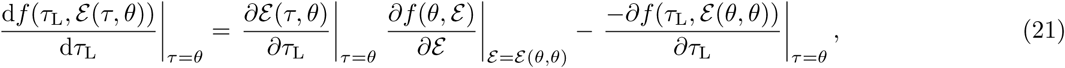

is directly influenced by the marginal cost of learning. Previous theoretical works have assumed that the marginal cost of learning is constant (and usually equals *ε* (*θ, θ*), see eq. 1 of Lehmann et al., 2013; eq. 11 of Wakano and Miura, 2014; eq. 4 of Kobayashi et al., 2015, eq. 4 of Ohtsuki et al., 2017, see Nakahashi, 2010 for an exception). Here, we do not make any such assumption, which allows us to evaluate the effects of different types of learning-reproduction trade-offs. Note that because cultural information *ε* is adaptive, we assume that fecundity is monotonically increasing with *ε* (i.e., *∂f* (*θ, ε*)*/∂ε >* 0, which is the “marginal benefit of information”). This means that the total derivative of fecundity with respect to learning *τ*_L_ (eq. 21) is proportional to

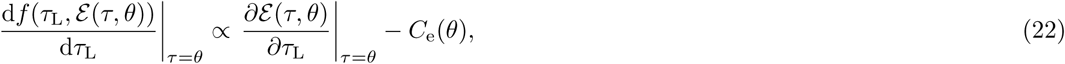

where

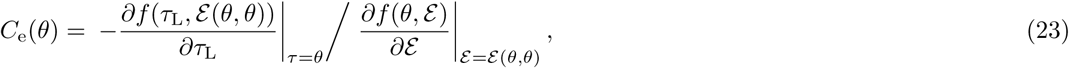

is the ratio of the marginal cost of learning to the marginal benefit of adaptive information. The quantity *C*_e_(*θ*) can be thought of as a marginal rate of substitution (Pindyck and Rubinfeld, 2001), i.e., how much an individual is ready to invest in learning in exchange for obtaining adaptive information while maintaining the same level of fecundity, which will turn out to be an important measure of learning cost. We refer to *C* _e_(*θ*) as the *effective* cost of learning.

### 2.3 Cultural information assumptions

#### 2.3.1 Cultural dynamics and resident cultural equilibrium

We now introduce the dynamics of adaptive information *ε_t_*(*τ, θ*) held by a heterozygote mutant in demographic time period *t* at the time of reproduction (after SL and IL occurred). Individuals acquire adaptive information by performing SL from the parental generation by way of vertical transmission with probability *v* and oblique transmission with probability 1 − *v*, but we assume that the efficiency of SL is independent of whether transmission occurs vertically or obliquely (see Kobayashi et al., 2015, for similar assumptions under haploid reproduction). The main dynamic assumption we make about adaptive information is that it satisfies the recursion

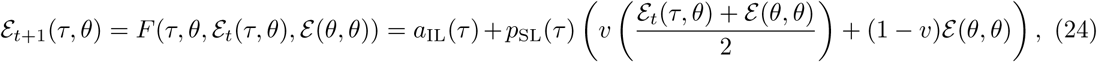

which depends on two terms. First, it depends on the information *a*_IL_(*τ*) an individual can obtain by performing IL alone. This is assumed to be equal to zero in the absence of effort *τ*_L_*τ*_IL_ devoted to IL, to increase and eventually saturate with effort *τ*_L_*τ*_IL_. Hence, *a*_IL_(*·*) is a function of a single argument: *a*_IL_(*τ*) = *a*_IL_(*τ*_L_*τ*_IL_), but we write it in terms of *τ* for ease of presentation. Second, adaptive information depends on the fraction *p*_SL_(*τ*) of information an individual obtains by SL from the parental generation. This is assumed to increase monotonically with the effort *τ*_L_(1 *− τ*_IL_) allocated to SL. So, *p*_SL_(*·*) is also a function of a single argument:*p*_SL_(*τ*) = *p*_SL_(*τ*_L_(1 *− τ*_IL_)).

The information obtained from the parental generation depends on the type of exemplar individual from which information is obtained. The interpretation of eq. (24) is that with probability *v* the cultural parent of the mutant is one of its two genetic parent, in which case with probability 1/2 the exemplar is a heterozygote mutant who carries information level *ε_t_*(*τ, θ*) and with probability 1/2, a homozygote for the resident who carries resident information level *ε*(*θ, θ*). With probability 1 *− v* the mutant performs oblique transmission, in which case the exemplar is a resident with the information level *ε*(*θ, θ*). The equilibrium resident level of information satisfies

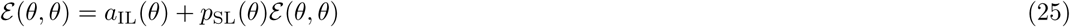

(by setting *τ* = *θ* in eq. 24 and letting *ε* (*θ, θ*) = lim_*t*→∞_ *ε_t_*(*θ, θ*)), or equivalently,

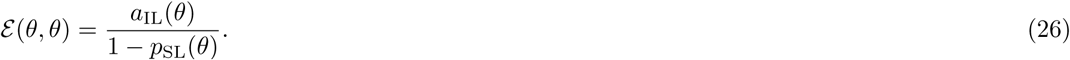

Our formulation of cultural dynamics in terms of IL and SL components (*a*_IL_(*τ*) and *p*_SL_(*τ*)) allows us to capture a variety of learning models that have previously been considered in the gene-culture co-evolution literature. For instance, eq. (24) allows to capture the classical models of IL and SL learning, where *ε* represents the probability of expressing the “correct” (or “optimal”) learned phenotype (e.g., light a fire), when an individual can express two alternative behaviors: the “correct” or the “wrong” phenotype (e.g., Rogers, 1988; Enquist et al., 2007; Kobayashi and Wakano, 2012; Aoki and Feldman, 2014; Wakano and Miura, 2014 and see section 2.3.2 for a concrete example). Equation (24) also allows to capture situations where the amount of information represents the total stock of knowledge or skill of an individual (e.g., Nakahashi, 2010; Aoki et al., 2012; Kobayashi and Aoki, 2012; Lehmann et al., 2013; Wakano and Miura, 2014; Kobayashi et al., 2015; Ohtsuki et al., 2017). Regardless of the precise cultural trait followed, eq. (24) embodies the feature that an individual can add up information by SL to that acquired by IL, which can result in a larger amount of cultural information at steady state, i.e., SL increases the amount of adaptive information by a factor 1/(1 *− p*_SL_((1 − *θ*_IL_)*θ*_L_)) (eq. 26). Hence, cumulative cultural evolution can occur.

#### 2.3.2 Mutant cultural equilibrium

From eq. (24), the mutant equilibrium level of cultural information is

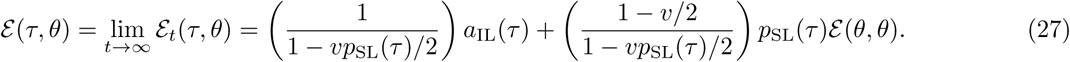

Comparing eq. (27) to eq. (25) shows that for a mutant who invests more in learning than residents (*a*_IL_(*τ*) *> a*_IL_(*θ*), *p*_SL_(*τ*) *> p*_SL_(*θ*)), vertical transmission (*v >* 0) increases both the level of individually and socially learned information relative to the baselines of *a*_IL_(*θ*) and *p*_SL_(*θ*) *ε*(*θ, θ*) obtained by a resident individual (eq. 25). Hence, vertical transmission increases the effective amount of IL and SL under selection.

### 2.4 Co-evolutionary equilibrium

We substitute the mutant information equilibrium (eq. 27) into the fitness function (eq. 20) to compute the selection gradients on the two evolving traits,

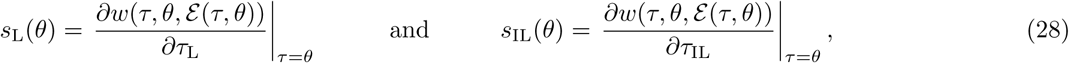

which describe the adaptive dynamics of the phenotypic traits that determine IL and SL. A necessary first-order condition for the learning rules to be locally uninvadable is then that *s*_L_(*θ\**) = 0 and *s*_IL_(*θ\**) = 0, where the associated cultural information *ε*(*θ, θ*) satisfies eq. (26). Whether the so-obtained singular strategies are convergence stable and locally uninvadable can be determined using the Jacobian and Hessian matrices eqs. (18)–(19) (see Appendix B).

We find that the selection gradient on learning can be expressed as

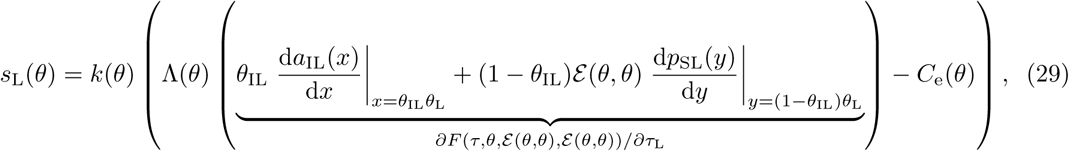

where

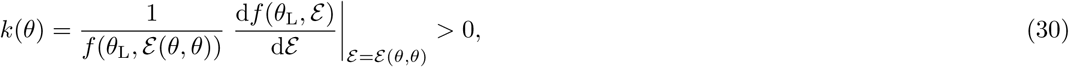

is a proportionality factor that does not affect the direction of selection,

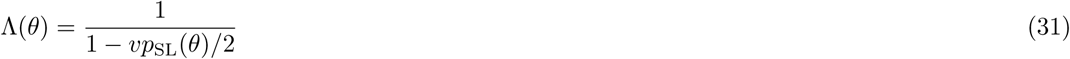

captures the multi-generational effects that relatives, including the focal, have on the information *ε* held by a focal individual (eq. 17), and *C*_e_(*θ*) is the effective cost of learning (eq. 23). Eq. (29) shows that the selection pressure on learning *θ*_L_ depends on (1) the marginal gain in information of a mutant (*∂F*(*τ, θ, ε*(*θ, θ*)*, ε*(*θ, θ*))*/∂τ*_L_, which is the average information gain from IL weighted by *θ*_IL_ and SL weighted by 1 − *θ*_IL_), due to the cumulated effects Λ(*θ*) of its own learning (“direct effect”) and that of its ancestors (“indirect effect”, see eq. 17); (2) the effective cost *C* _e_(*θ*) of learning (eq. 23).

Meanwhile, we find that the selection gradient on IL (*θ*_IL_),

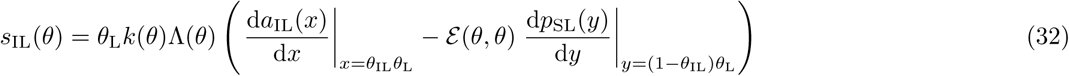

balances the marginal benefit of IL (d*a*_IL_(*x*)/d*x*) and the marginal cost (− d*p*_SL_(*y*)/d*y*) of allocating less resources to SL. Interestingly, vertical transmission does not directly influence the direction of selection on IL. Its influence may however be indirect through it effects on *θ*_L_ and therefore on the marginal benefits and cost of IL.

The selection gradient on learning (eqs. 29–31) shows that vertical transmission, by increasing the multi-generational effects Λ(*θ*) (eq. 31), amplifies the benefits of learning (eq. 29). This can be understood by considering that under vertical transmission, gathering adaptive information will later preferentially benefit relatives. But since *p*_SL_(*y*) *<* 1, vertical transmission can amplify the benefits of learning by at most a factor of 2 in this model (when *v* = 1, eq. 31). This moderate impact of vertical transmission is due to the assumptions underlying information dynamics (eq. 24): individuals are diploids, vertical transmission is equally likely to occur between a mutant offspring and its mutant and resident parents, and vertical transmission has no effect on the efficiency of transmission. As a result of these assumptions, information transformation eq. (24) depends linearly on vertical transmission rate *v* and only by a factor of 1/2, which limits the effect of vertical transmission.

More generally, the effect of vertical transmission on the amplification factor of learning benefits is given by

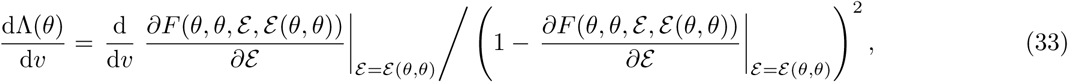

which depends on the effect of *v* on how the mutant environment affect its own dynamic. Eq. (33) reveals that if vertical information has a large effect on cultural dynamics, then it can have a large impact on Λ(*θ*), and thus have a significant influence on the evolution of learning strategies (eq. 11) and cumulative culture. This will occur for instance if for the same effort allocated to SL, the fraction of information *p*_SL_ obtained socially from an exemplar depends linearly on *v* (e.g., if offspring learn at a faster rate from their genetic parent than from unrelated individuals), or if *ε* depends non-linearly on *v* (e.g., if offspring first learn from their genetic parent and then learn obliquely at a different rate). The assumptions of our model (eq. 24) are therefore conservative for the effect of vertical transmission on cumulative culture. We endorsed them because they allow to make direct comparison with models in the literature, to obtain manageable analytical expression, and to check the validity of our results with individual-based simulations, three endeavours to which we next turn.

### 2.5 Evolution of learning to obtain the “correct” phenotype

In order to fully work out a concrete application of our approach, we make some further assumptions about the dynamics of the cultural information, and let *ε* be the probability that at the stage of reproduction an individual expresses the “correct” phenotype in its environment (e.g., Rogers, 1988; Enquist et al., 2007; Aoki and Feldman, 2014; Wakano and Miura, 2014). We assume that learning occurs on a fast time scale that is embedded in a single demographic time period (see Appendix C for details). In this mechanistic model, individuals first perform SL from the parental generation for (1 *− τ*_IL_)*τ*_L_ units of time (on the fast time scale) at rate *β*, proportionally to the distance between the probabilities *ε* of the exemplar and target individuals. Individuals then perform IL for *τ*_IL_*τ*_L_ units of time at a rate proportionally to the distance between its current probability *ε* and the upper bound 1 (since it is a probability). In this case, we show in Appendix C that the cultural level obtained by an individual can be written in the form

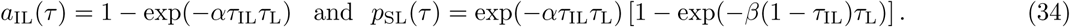

Interestingly, eq. (34) also captures the “critical social learning strategy”, where an individual first learns from the parental generation by SL and then, if it has not acquired the correct phenotype, performs IL (Enquist et al., 2007; Rendell et al., 2010, see also eq. (C-6) in Appendix C for the interpretation of eq. 34 in terms of the critical social learner strategy).

We assume that fecundity depends on whether the correct phenotype is acquired, and write it as

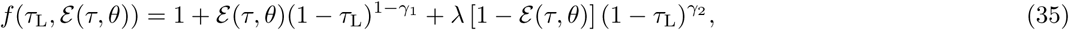

where “1” is the baseline reproductive unit and the rest can be understood as follows. A mutant has (1 *− τ*_L_) units of resources left to allocate into reproduction and the effect of this allocation on fecundity depends on whether a mutant expresses the correct phenotype. When the mutant has the “correct” phenotype (with probability *ε* (*τ, θ*)), returns on investment are controlled by *γ*_1_ and we assume that 0 ≤ *γ*_1_ *<* 1 so that returns rise linearly (*γ*_1_ = 0) or sharply with investment (*γ*_1_ *>* 0, Fig. 1). When the mutant has the “wrong” phenotype (with probability 1 *− ε*(*τ, θ*)), returns are tuned by *γ*_2_ and we assume that *γ*_2_ *>* 1 so that returns increase slowly with initial investment. Finally, the parameter 0 ≤ *λ* ≤ 1 bounds the fecundity of an individual with the wrong phenotype (Fig. 1). For example, an individual with the wrong phenotype who invests all resources into reproduction (*τ*_L_ = 0) has a fecundity 1 + *λ*. The parameters *γ*_1_, *γ*_2_ and *λ* in eq. (35) therefore influence how resources allocated into learning trade-off with fecundity.

**Figure 1:**
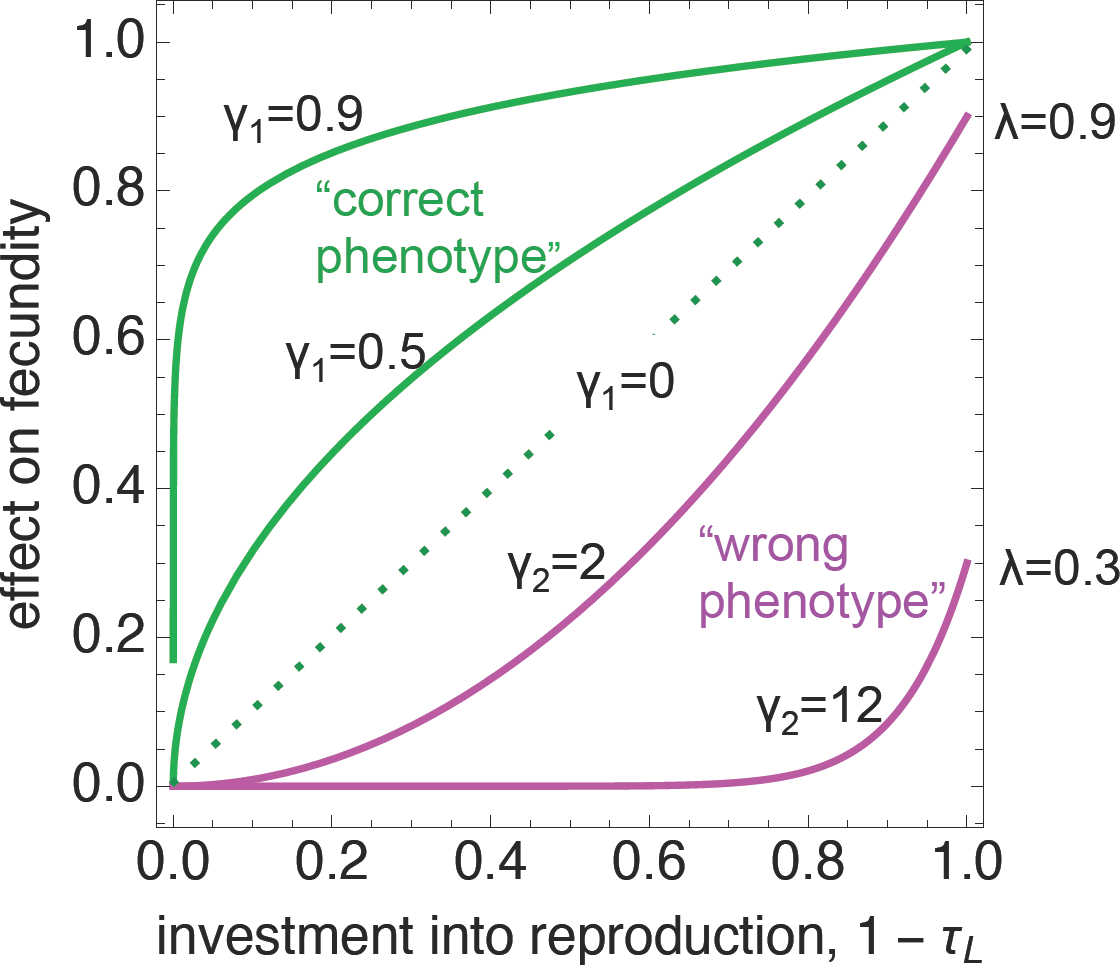
Components of the fecundity function (eq. (35)) as a result of expressing the correct or wrong phenotype. When a mutant expresses the correct phenotype, it obtains rapidly rising returns on investment ((1 *− τ*_L_)^1 *−γ*_1_^ where 0 ≤ *γ*_1_ < 1 tunes the sharpness of the rise, in green). When a mutant expresses the wrong phenotype, it obtains slowly rising returns on investment (*λ*(1 *− τ*_L_)^*γ*2^ where *γ*_2_ *>* 1 controls how slowly the returns are and 0 ≤ < 1 bounds the returns, in pink). The total effect on fecundity eq. (35) is then given by the average returns, averaged over the probabilities of expressing the correct (*ε*(*τ, θ*)) and wrong (1 *− ε*(*τ, θ*)) phenotype.

#### 2.5.1 Effect of vertical transmission on cumulative culture

We first work out the case where expressing the wrong phenotype results in zero effects on fitness (*λ* = 0, the usual case in the literature, Rogers, 1988; Enquist et al., 2007; Kobayashi and Wakano, 2012; Wakano and Miura, 2014; Aoki and Feldman, 2014), for which singular learning strategies 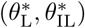 and the corresponding levels of information *ε\** they generate can be determined analytically. Substituting eqs. (34)–(35) into eq. (32), we find that solving for 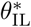 such that 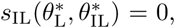 there is a unique IL singular strategy

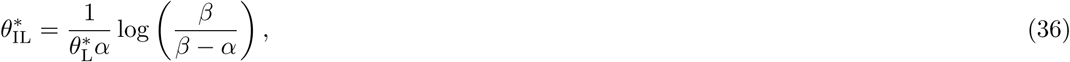

which rapidly (hyperbolically) decreases with the resources 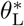 allocated to learning, but is otherwise independent of the rate of vertical transmission (as predicted by eq. 32). Eq. (36) also shows that for individuals to evolve a composite learning strategy 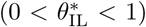 that mixes SL and IL, it is necessary (but not sufficient) that the rate of SL is greater than the rate of IL (*β > α*), otherwise SL cannot be a singular strategy 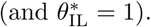

Substituting eqs. (34)–(35) into eq. (29) with *λ* = 0, we find that learning increases from zero (when *θ*_L_ = 0 and *θ*_IL_ = 1) when *>* 0 (i.e., *s*_L_(0, 1) *>* 0 when *α >* 0). Then, solving 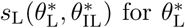 such that 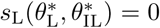 using eq. (36), we find that there is a unique singular level of learning that is given by

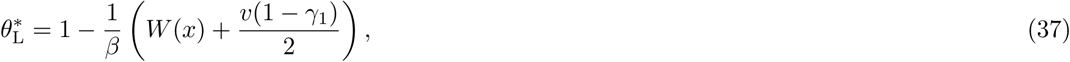

where *W*(*x*) is the principal solution of the Lambert function (the solution for *y* in *x* =*ye^y^*) with argument

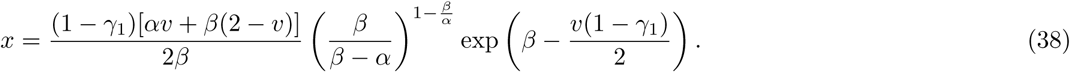

Then, by substituting the singular strategies eqs. (36)-(37) into eq. (26), we find that the singular interior learning strategy generates a level

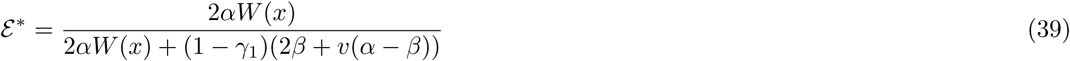

of adaptive information. If, like Enquist et al. (2007), we assume that learning has no marginal cost (i.e., *γ*_1_ = 1), that all individuals engage into learning (*θ*_L_ = 1), and that there is no vertical transmission (*v* = 0), we recover Enquist et al. (2007)’s result: efficient social transmission of information (*β* ≫ 0) promotes high levels of adaptive information (see Appendix D, eqs. D-1-D-3 for details).

More generally, however, a numerical inspection of the singular learning strategies 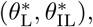 and the corresponding amount of adaptive information *ε\** they generate, highlight the difficulty for adaptive information to accumulate (Fig. 2). In particular, even though vertical transmission *v* increases the amount of time invested into learning and adaptive information, its effect is moderate (Fig. 2, top row). In fact, from eq. (39), we see that under pure oblique transmission (*v* = 0), the level of information converges asymptotically to

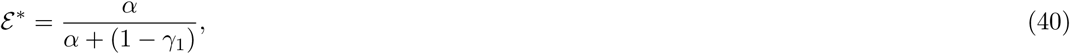

as *β → ∞*, while under pure vertical transmission (*v* = 1), it converges to

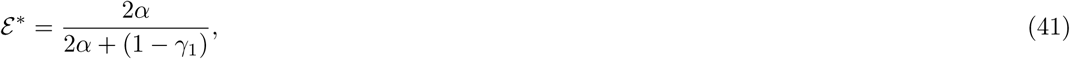

which shows that vertical transmission cannot more than double the amount of information. This is in line with our result that when vertical transmission has no effect on the efficiency of transmission and the dynamic dependence of *ε* is linear in *v* (eq. 31), transmission has a moderate effect on cumulative cultural evolution (section 2.4).

**Figure 2:**
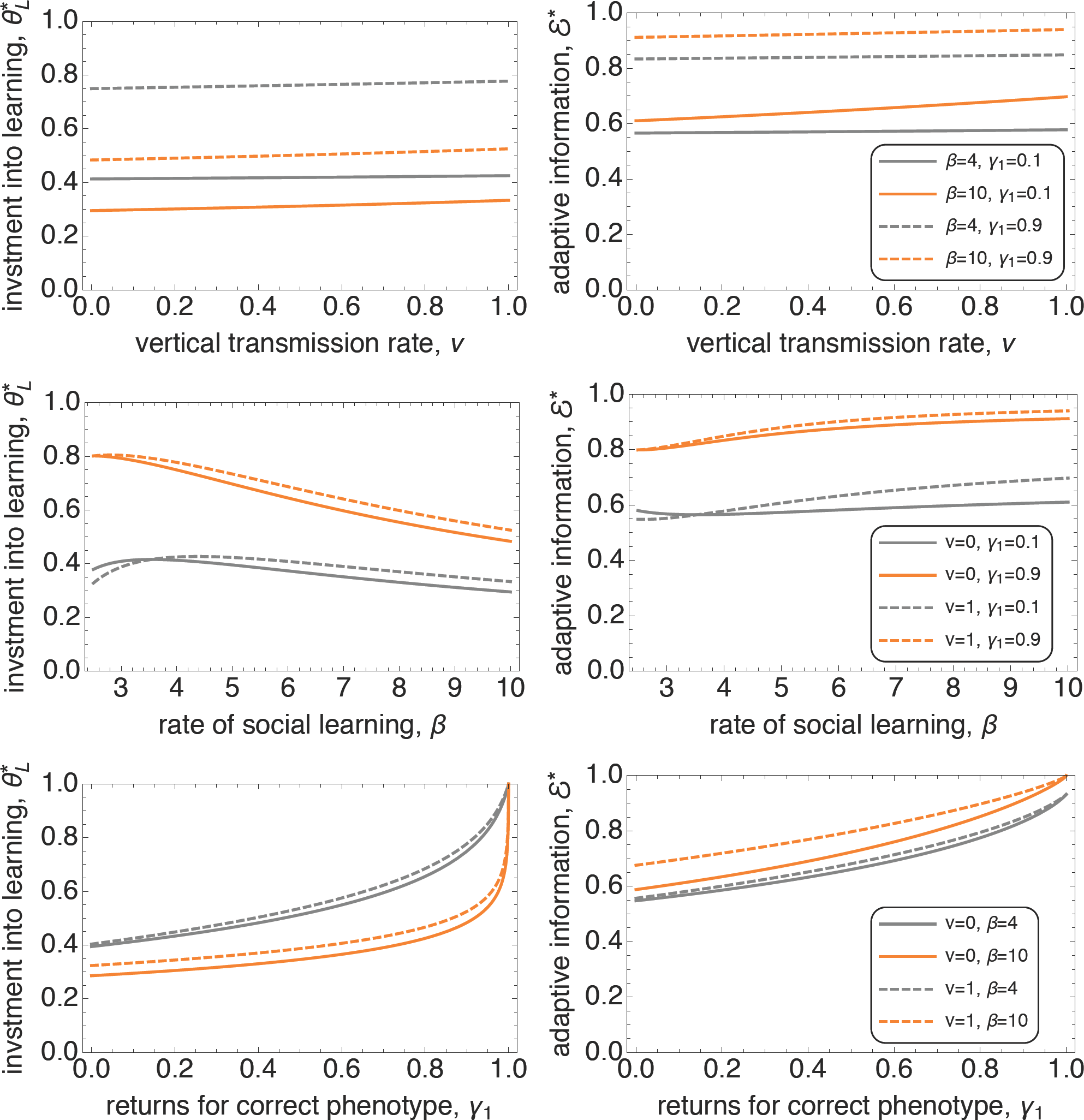
Singular learning strategy 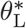 and the corresponding level of adaptive information *ε\** it generates (obtained from eqs. (36)-(39)) as a function of the vertical transmission rate *v* (Top row), the rate of social learning *β* (middle row) and the returns for expressing the correct phenotype *γ*_1_ (bottom row). Different lines (gray, orange, full and dashed) stand for different parameter values, whose values are shown in the inset of each row (rate of individual learning is fixed *α* = 2 everywhere).

Our finding that vertical transmission has little effect on cumulative cultural evolution is also consistent with the results of Kobayashi et al. (2015), who considered a haploid population when adaptive information is the total stock of knowledge held by an individual at the stage of reproduction and marginal learning costs are constant (with *λ* = *γ*_1_ = 0, see Appendix D eqs. D-4–D-11 for an application of our framework to Kobayashi et al., 2015’s model). A comparison with Kobayashi et al. (2015)’s results allows us to evaluate the role of diploidy on cumulative culture when marginal learning costs are constant (*γ*_1_ = 0). If Kobayashi et al. (2015)’s model showed a limited effect of intermediate vertical transmission (0 *< v <* 1), it showed that pure vertical transmission (*v* = 1) could have a strong positive influence on learning. This strong effect arises because when *v* = 1, the amplification factor Λ(*θ*) of learning benefits can go to infinity (see eq. D-11 in Appendix D). By contrast, the effect of pure vertical transmission (*v* = 1) remains moderate in diploids because relatedness between a focal individual and its exemplar is half of that under haploid reproduction and so the amplification factor Λ(*θ*) is at most 2 (eqs. 29 and 31).

The other parameters of the model also affect the singular learning strategies 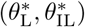 and therefore the level of adaptive information *ε\** at equilibrium. The rate of SL *β* has a negative effect on the level of 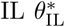 (eq. 36), and hence positive effect on SL, but only a moderate effect on adaptive information *ε\** (Fig. 2, middle row). By contrast, the returns of investment into reproduction *γ*_1_ when expressing the correct phenotype significantly increase both the investment into learning 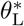o and adaptive information *ε\** (Fig. 2, bottom row). This is because *γ*_1_ decreases the marginal cost of learning,

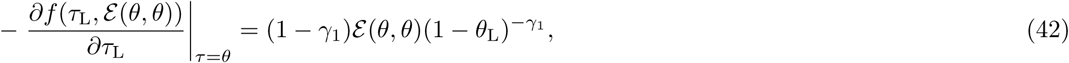

and hence the effective cost of learning,

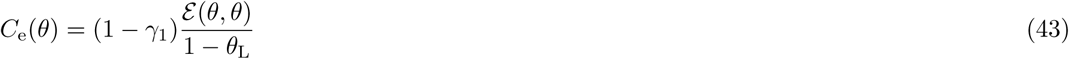

Previous studies have in their majority assumed that marginal learning costs are constant (*γ*_1_ = 0 so that the marginal cost is *ε*(*θ, θ*)), which results in high effective learning cost (*C*_e_(*θ*) = *ε*(*θ, θ*)/(1 − *θ*_L_), Lehmann et al., 2013; Wakano and Miura, 2014; Kobayashi et al., 2015; Ohtsuki et al., 2017). Our results therefore highlight the importance of the marginal and effective cost of learning, or more generally of the learning-reproduction trade-off. Note that for all parameter values displayed in Fig. 2, we checked numerically (by applying eqs. 18–19, see Appendix B) that singular learning strategy 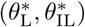 defined by eqs. (36)–(39) is convergence stable (Fig. 3) and locally uninvadable, which was confirmed by individual based simulations (Fig. 4).

**Figure 3:**
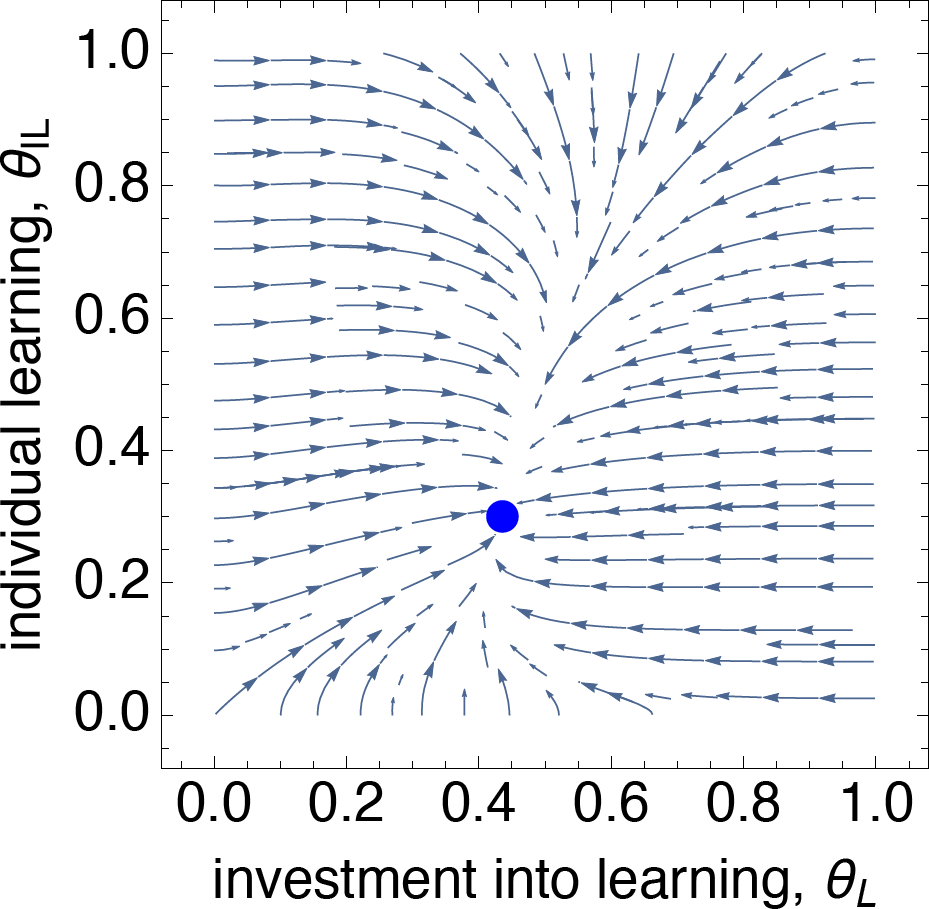
Phase portrait of the co-evolutionary dynamics of *θ*_L_ and *θ*_IL_ given by the selection gradients on each corresponding trait (eqs. 29 and 32 using eqs. 34–35 and *λ* = 0, other parameters: *α* = 1, *β* = 8, and *γ*_1_ = 0.5 and *v* = 0.75). The evolutionary dynamics converge to the single interior point shown in blue 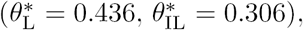 which is convergence stable as can be seen from the graph, and which can also be shown to be locally uninvadable (using Appendix B).

**Figure 4:**
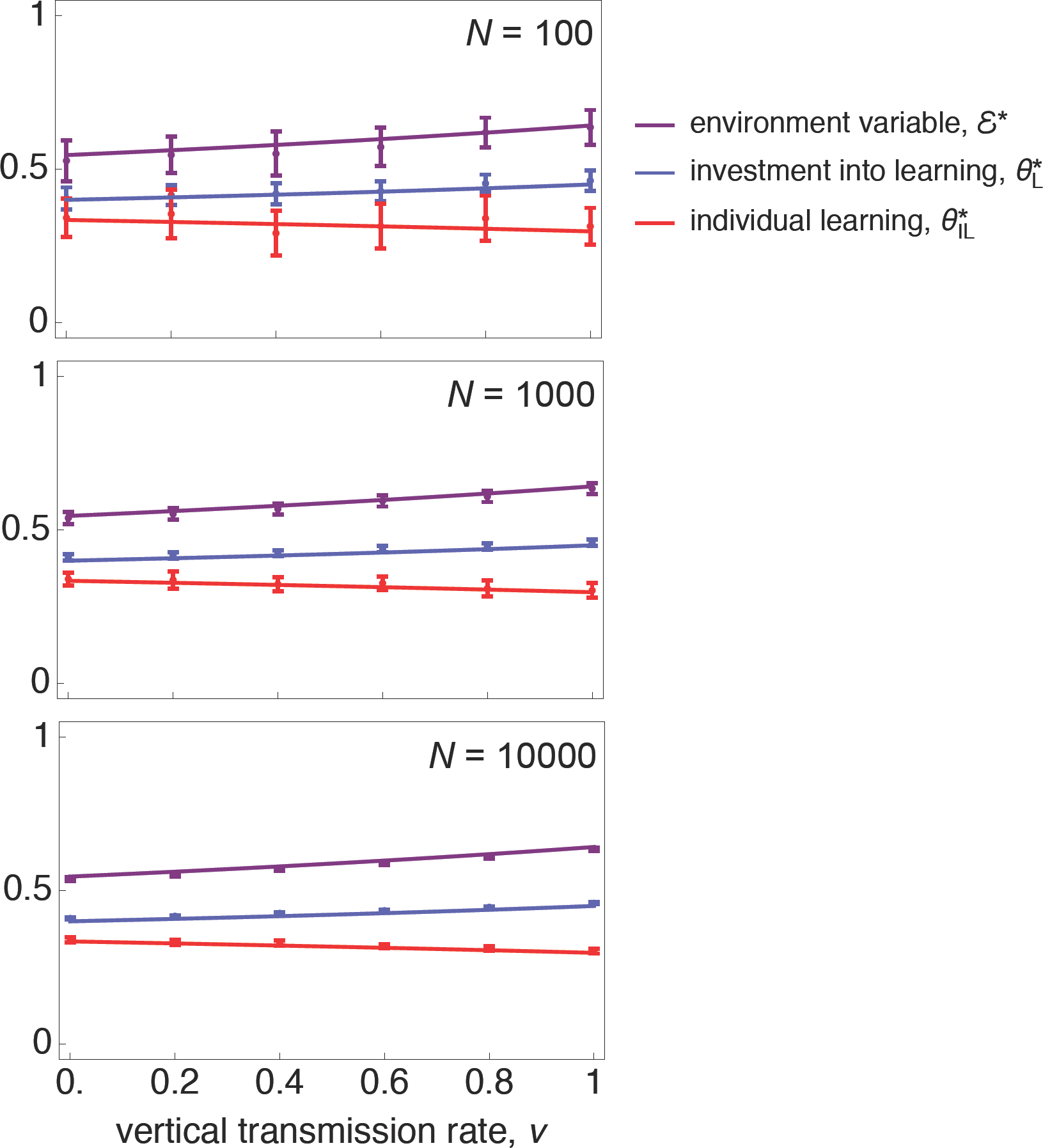
Comparison between analytical results and individual-based simulations. In each panel, analytical equilibria are shown in full lines 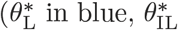 in red, and *ε\** in purple given by eqs. 36-39, with *α* = 1, *β* = 8, and *γ*_1_ = 0.5). The results of simulations (described in Appendix E) are given by the temporal population trait averages (points, same colour as lines) and the temporal standard deviation of population trait averages (error bars centred on averages, same colour as lines). The top panel is for a population size of *N* = 100, middle for *N* = 1000, and bottom for *N* = 10000, which show a good agreement between analytical predictions and simulation results, especially for large populations in which genetic drift has little effect and in which deviations between invasion analysis results and exact results for finite population based on fixation probabilities are of the order 1 */N* (Rousset, 2004).

#### 2.5.2 Effect of vertical transmission on evolutionary branching

We now turn to the case in which individuals with the wrong phenotype can obtain returns on investment in fecundity, *λ >* 0, and consider that an individual with the correct phenotype does not need to invest much into reproduction to obtain a fecundity gain (concave returns, 0 *< γ*_1_ *<* 1), while an individual with the wrong phenotype needs to invest significantly more into reproduction to have a fecundity gain (convex returns, *γ*_2_ *>* 1, see Fig. 1).

We find that the singular strategy for individual learning *θ*_IL_ is the same as when *λ* = 0 (eq. 36), but we are unable to obtain a general analytical solution for singular strategies for investment into learning (*θ*_L_). The model is therefore studied numerically and the main qualitative outcomes are as follows (see Appendix F for details on the numerical approach). By contrast to our results with *λ* = 0, it is more difficult for learning (*θ*_L_) to evolve from zero and the evolutionary dynamics to converge to an interior equilibrium 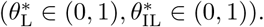. This is because when individuals with the wrong phenotype obtain a significant fecundity benefit when all resources are invested into reproduction (*θ*_L_ = 0), a mutant that invests only a little into learning cannot compete with a resident who invests all into reproduction. Only over a critical threshold of learning does extra investment into learning become beneficial compared to the cost of diverting resources from reproduction, in which case both learning and SL can be favored (see Fig. 5).

**Figure 5:**
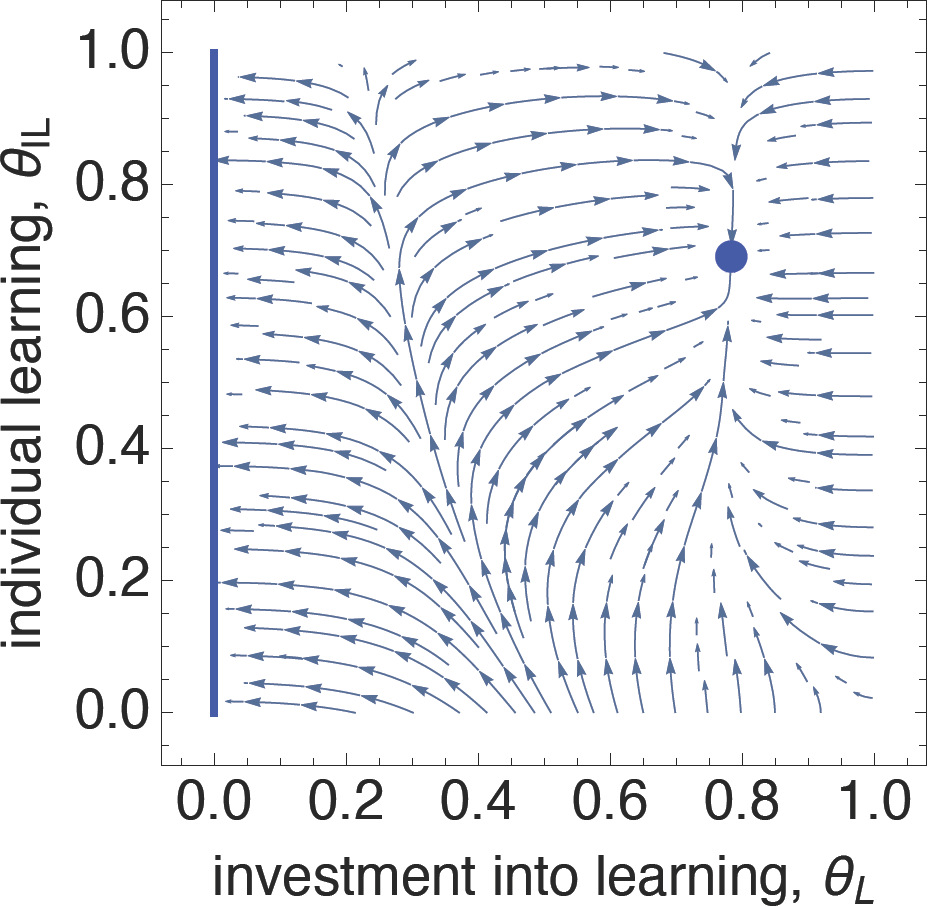
Phase portrait given by the selection gradients on each trait (eqs. 29 and 32 with *v* = 0, *α* = 2, *β* = 3, *γ*_1_ = 0.9, *γ*_2_ = 3, = 0.7) and evolutionary convergence equilibria shown in blue (blue line for zero, and filled blue circle for convergence stable interior point), which shows that either *θ*_L_ converges to zero, or both *θ*_L_ and *θ*_IL_ converge to an interior equilibrium 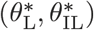 depending on the initial population values. As the graph shows, the co-evolutionary dynamics of *θ*_L_ and *θ*_IL_ exhibit bistability and the population must cross a threshold level of learning in order for selection to favor greater investment into learning.

When a mixed convergence stable interior strategy exists 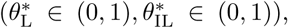, our analysis suggests that it is usually uninvadable (Appendix F). We found nevertheless parameter values for which convergence stable interior equilibria are locally invadable (i.e., disruptive selection occurs), so that evolutionary branching (Geritz et al., 1997, 1998) can happent at these. Simulations showed that when it occurs, evolutionary branching leads to the emergence of two morphs, one that does less IL than the equilibrium, and who selfishly exploits the other that does more IL than the convergence stable equilibrium (Fig. 6). We find that vertical transmission is important for disruptive selection, in general inhibiting it (Appendix F and Supplementary Figure 1). This is because vertical transmission prevents adaptive cultural information from being transmitted from lineages that perform individual learning to others that do not. Our finding that vertical transmission, which increases the correlation among genes and culture, inhibits disruptive is in line with the finding that a reduction in migration (which increases relatedness) inhibits disruptive selection in spatially structured populations (Ajar, 2003; Mullon et al., 2016).

**Figure 6:**
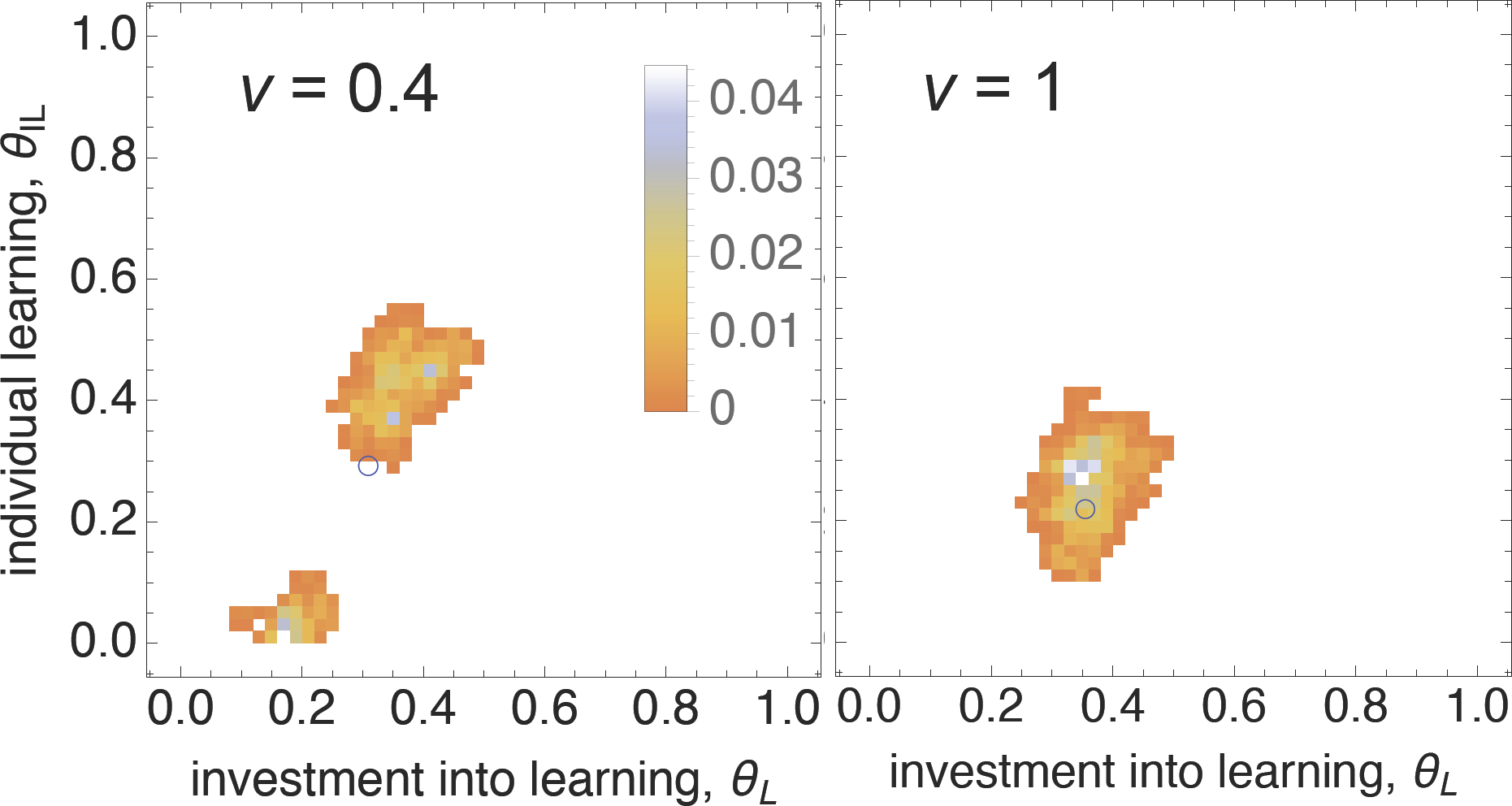
Evolutionary branching and stability. Bivariate distributions of trait values (*θ*_L_ and *θ*_IL_), depicted as two-dimensional colour histograms with bins of width 0.02 and in which bin colour gives the frequency of individuals within each bin (see figure legend), obtained under individual-based simulations (frequencies averaged over the last 10^3^ generations after 9.9 *×* 10^4^ generations of evolution, parameters: *α* = 2, *β* = 12, *γ*_1_ = 0.7, *γ*_2_ = 5, *λ* = 0.7, Appendix E for details). When *v* = 0.4, the population is split into two morphs around the interior convergence stable equilibrium 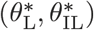 (blue empty circle), but when *v* = 1 the population is unimodal, centred around the convergence stable equilibrium (blue empty circle).

## 3 Discussion

In spite of the evidence that cultural transmission often occurs between parent and offspring (Cavalli-Sforza and Feldman, 1981; Guglielmino et al., 1995; Hewlett et al., 2011; Konner, 2010), the bulk of theoretical work on gene-culture co-evolution for cumulative culture has focused on oblique or horizontal transmission (e.g., Boyd and Richerson, 1985; Rogers, 1988; Enquist et al., 2007; Rendell et al., 2010; Nakahashi, 2010; Aoki et al., 2012; Lehmann et al., 2013; Nakahashi, 2013; Aoki and Feldman, 2014; Wakano and Miura, 2014). In this paper, we derived the invasion fitness of a mutant allele that co-evolves with cumulative cultural information under vertical transmission. We showed that when cultural dynamics are deterministic and have a single attractor equilibrium, invasion fitness is equal to the *individual fitness* of a mutant when the dynamic of cultural information is at equilibrium (eq. 5). This result allows for an analytically tractable study of gene-culture co-evolutionary adaptive dynamics in family-structured populations. Our invasion fitness measure (eq. 5) can also be applied to cases where population size varies, following a deterministic dynamic with a single attractor point (in which case fitness eq. 7 is evaluated at this attractor for the resident population, e.g., Ferrière and Gatto, 1995; Caswell, 2000; Metz, 2011).

Our analysis of the selection gradient revealed that cultural evolution results in carry-over effects across generations, which feedback on the selection pressure of traits affecting cultural dynamics (see eq. 15). This feedback arises from kin selection, since an individual modifying cultural information as a result of expressing a mutant allele will change the environment to which descendants living in the next or more distant generations are exposed. These indirect fitness effects depend on the magnitude of how altering trait values affect cultural information dynamics and how the resulting change in dynamics affects individuals in downstream generations (eq. 15). From the perspective of a recipient of these effects, its fitness will depend on a multitude of individuals in past generations (all those from the ancestral lineage of an individual carrying a mutation). These multi-generational effects accumulate in a geometric progression over the lineage of a mutant and can therefore potentially be large (see eq. 17). More broadly, our approach allows for a clear separation between direct and indirect effects on fitness and can be readily extended to cover more realistic demographic scenarios such as aged-structured populations with senescence.

We applied our framework to a generic model in which genetically determined phenotypes that underlie individual learning (IL) and social learning (SL) co-evolve with the adaptive information they generate (eq. 24), and which generalizes many previous scenarios for the co-evolution of IL and SL (Rogers, 1988; Enquist et al., 2007; Aoki and Feldman, 2014; Wakano and Miura, 2014; Nakahashi, 2010; Aoki et al., 2012; Lehmann et al., 2013; Wakano and Miura, 2014; Kobayashi et al., 2015). The analysis of the selection gradient showed that the impact of vertical transmission, which increases interactions between relatives and thus kin selection effects, can be viewed as an amplification factor of the benefits of learning (eqs. 29 and 31). Vertical transmission therefore favors greater levels of learning and hence cumulative culture. But when transmission is as efficient between relatives than between unrelated individuals, learning benefits cannot be amplified by more than a factor of two in randomly-mating diploids (eq. 31). By contrast, in haploids or when transmission is only among heterozygotic mutant parents and offspring, learning benefits can explode under pure vertical transmission (*v* = 1, eq. D-11).

The selection pressure on phenotypic carry-over effects impacting relatives is therefore diluted in the presence of diploidy. This is because diploidy decreases the correlation between genetic and cultural transmission pathways so that over generations, mutant alleles also benefit resident ones. The correlation between genes and vertically transmitted cultural information could be enhanced for instance by assortative mating between genetically-similar individuals, which is not taken into account by our formalization, and which would presumably offset the dilution effect of diploidy we observe. Assortative mating, whether it occurs among genetically or culturally similar individuals, has been shown to influence gene-culture coevolution when selection is frequency-independent (Creanza et al., 2012), in which case culture cannot be cumulative. In the future, it would be relevant to investigate the impact of assortative mating on cumulative cultural evolution and its interaction with vertical transmission, as it may magnify selection on multigenerational effects.

In order to illustrate our results, we analyzed in detail a specific scenario of the evolution of IL and SL where adaptive information determines the acquisition of a binary trait describing “correct” and “wrong” phenotype to be expressed in a given environment (e.g., light a fire, see eq. (34) and Rogers, 1988; Wakano et al., 2004; Wakano and Aoki, 2006; Enquist et al., 2007; Rendell et al., 2010; Kobayashi and Wakano, 2012; Wakano and Miura, 2014), and also provided a mechanistic derivation of the critical social learner strategy (Enquist et al., 2007; Rendell et al., 2010, see Appendix C). We found that in the absence of vertical transmission (*v* = 0) and with constant marginal learning costs (*γ*_1_ = 0 and = 0), it is difficult for culture to accumulate. This corroborates Wakano and Miura (2014)’s conjecture that in general, learning-reproduction trade-offs make it more difficult for cumulative culture to evolve.

When transmission occurs vertically, kin selection has a positive yet moderate quantitative impact on the amount of cultural information accumulated by individuals in uninvadable populations for arbitrary marginal learning costs (Fig. 2, Top row). This generalizes the results of Kobayashi et al. (2015), who studied a haploid model with constant marginal learning costs. The weak kin selection effects on cumulative culture we observe also echo Ohtsuki et al. (2017)’s results on the evolution of IL and SL, when cultural transmission occurs within spatial groups connected by limited dispersal. In this case, oblique transmission occurs between genetically related individuals, which increases positive kin selection effects on cumulative culture. But limited dispersal also increases kin competition, which disfavors the transmission of adaptive cultural information. Overall, Ohtsuki et al. (2017) found that kin selection has an appreciable effect on cumulative culture only under extreme kin structure (small local groups, low dispersal, which is akin to markedly increase Λ(*θ*) in eq. 29). Our results are therefore in line with the notion that kin selection has weak effects on the accumulation of adaptive information across generations, at least when information is transmitted as efficiently between family members as between unrelated individuals.

By contrast, we showed that the parameter *γ*_1_, which influences the costs of learning (eq. 42-43), can have a significant quantitative influence on the uninvadable level of adaptive information and affect cumulative culture by an order of magnitude (Fig. 2, bottom row). Our results therefore stress the importance of the shape of the learning-reproduction trade-off, which has perhaps been under-appreciated by a literature that has mostly considered constant marginal learning costs (e.g., Rogers, 1988; Wakano et al., 2004; Wakano and Aoki, 2006; Enquist et al., 2007; Rendell et al., 2010; Kobayashi and Wakano, 2012; Lehmann et al., 2013; Kobayashi et al., 2015; Ohtsuki et al., 2017, but see Nakahashi, 2010 for an exception). Like in our model, non-linear marginal learning costs will arise when expressing a correct cultural phenotype entails rapidly rising returns of resources allocated into reproduction (*γ*_1_ *>* 0, Fig. 1), which is a plausible biological scenario. For instance, being able to light a fire, which allows warming and feeding a high energy diet to one’s offspring, is likely to produce sharply rising fecundity returns. However, in the absence of any empirical quantification of learning costs and benefits, it is difficult to make any conclusion about their role relative to vertical transmission on cultural evolution in natural populations. The quantification of these costs and benefits across the lifespan of individuals unfortunately remains a neglected topic in cultural evolution (Demps et al., 2012).

To conclude, we have shown that when the effect of learning on the adaptive information held by an individual is independent from whether learning occurs between family members or between unrelated individuals, vertical transmission moderately increases levels of adaptive cultural information through kin selection effects. But our model suggests that if vertical transmission interacts with cultural dynamics, then vertical transmission may have a large influence on cumulative culture (eqs. 15 and 33). This would occur for example if offspring learn better from relatives, or if parents devote more teaching effort towards offspring than towards unrelated individuals. Such scenarios would be relevant to investigate in future research, and also raise the question of the evolution of vertical transmission itself. In addition, our finding that vertical transmission inhibits disruptive selection suggests that it can play a qualitative role in the evolution of IL and SL learning strategies themselves (e.g., conformist transmission, payoff biased transmission, teaching), whose evolution under vertical transmission have not been much investigated. Our approach can be readily accommodated to study all these specific questions, and more broadly, questions on niche construction and gene-culture co-evolution within the family where the modified rates of cultural transmission have long-lasting effects on future generations.

## Acknowledgments

We thank Yutaka Kobayashi for useful discussions about cultural evolution.

## Appendices

### Appendix A: invasion fitness

Given the definition of individual fitness *w*(*τ, θ, ε_t_*(*τ, θ*)) in the main text and the associated dynamics for the cultural information *ε_t_*(*τ, θ*) (eq. 2), our modelling assumptions entail that invasion fitness is given by the geometric growth rate

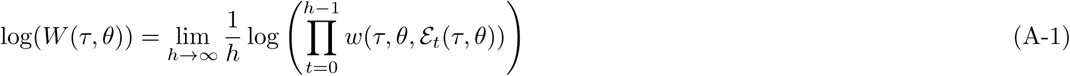

(Cohen, 1979; Tuljapurkar, 1989; Charlesworth, 1994; Ferrière and Gatto, 1995; Caswell, 2000; van Baalen, 2013). It is in general challenging to evaluate eq. (A-1) explicitly under an arbitrary dynamics of *ε_t_*(*τ, θ*) (Tuljapurkar, 1989). But since we assume that the sequence of environments 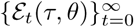 converges to a unique fixed point *ε*(*τ, θ*) (satisfying eq. 4), the limit of eq. (A-1) converges to

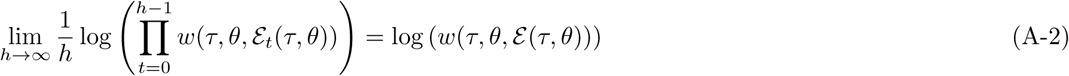

(see also Ferrière and Gatto, 1995), and so invasion fitness is given by

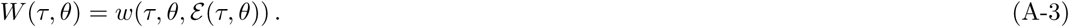

### Appendix B: second-order conditions

From eq. (18) we can evaluate the condition of strong convergence stability for a singular point = 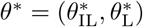 satisfying eq. (10). This requires that the Jacobian matrix

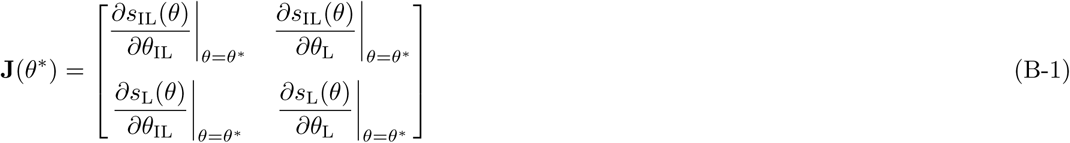

has eigenvalues with negative real parts (e.g., Lessard, 1990; Leimar, 2009; Mullon et al., 2016). Meanwhile, a singular point 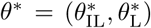 is uninvadable when the Hessian matrix

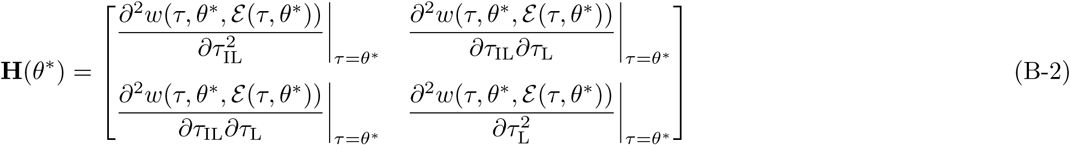

is negative-definite, or equivalently that both its eigenvalues are negative (note that since H(*θ\**) has real entries and is symmetric, its eigenvalues are necessarily real, e.g., Lessard, 1990; Mullon et al., 2016).

### Appendix C: cultural dynamics

We derive here eq. (34) by assuming a fast time scale of learning within a single demographic time period and by setting the total time length of learning to unity. This approach is equivalent to previous models of cultural information (Aoki et al., 2012; Lehmann et al., 2013; Wakano and Miura, 2014; Kobayashi et al., 2015; Ohtsuki et al., 2017).

We start by assuming that the resident population is at its equilibrium for cultural dynamics (satisfying eq. 3). Then, an individual first learns from the parental generation (either vertically or obliquely) at rate *β* per unit time and we assume that the rate of change of the adaptive information *ε*(*h*) held at time *h* of the fast time scale of an individual is

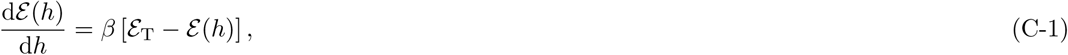

which for a mutant holds for *h ∊* [0, (1 − *τ*_IL_)*τ*_L_] (i.e., (1 *− τ*_IL_)*τ*_L_ is the time spent performing SL) and where *ε*_T_ is the cultural information of the exemplar individual (or cultural parent) and the initial condition is *ε* (0) = 0. Eq. (C-1) entails that an individual acquires the correct phenotype proportionally to *β* and the difference between the probability that the focal and the exemplar individual has the correct phenotype. The solution of eq. (C-1) is

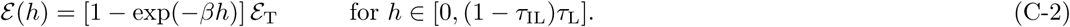

After SL has been performed, the individual performs IL and the rate of change in adaptive information ε(*h*) during IL is given by

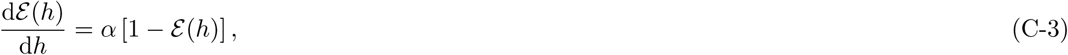

which for a mutant holds for *h ∊*((1 *− τ*_IL_)*τ*_L_*, τ*_L_] (i.e., *τ*_IL_*τ*_L_ is the time spent performing IL) with initial condition *ε* ((1 *− τ*_IL_)*τ*_L_)) (given by eq. C-2). According to eq. (C-3), the individual acquires the correct phenotype by IL proportionally to and its current distance to the “target” which is 1. This formulation implements standard reinforcement learning (Bush and Mosteller, 1951). The solution to eq. (C-3) is

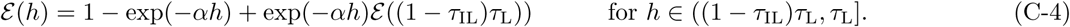

Combining eq. (C-2) and eq. (C-4) yields that at the final time of the learning period (*h* =*τ*_L_), say at generation *t*, the amount of adaptive information held by a mutant is

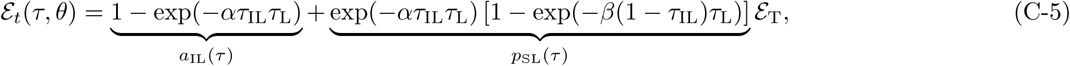

which yields the components of eq. (34) of the main text and where *ε*_T_ is equal to the average cultural information of the cases when the individual performs vertical transmission ((*ε_t−1_*(*τ, θ*) + *ε*(*θ, θ*))/2) and when it performs oblique transmission (*ε*(*θ, θ*)).

Interestingly, the right hand side of eq. (C-5) can be equivalently written as

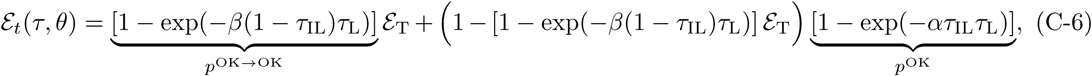

where *p*^OK→OK^ is the probability of obtaining the correct phenotype via social learning from an individual with the correct phenotype, and *p*^OK^ is the probability of obtaining the correct phenotype via individual learning. Eq. (C-6) shows that we can interpret this model in terms of an individual first attempting to learn the “correct” solution by SL, and if it is unsuccessful, it tries to acquire the correct phenotype by IL. This is equivalent to the learning strategy called *critical SL* (Enquist et al., 2007; Rendell et al., 2010).

### Appendix D: connection to previous work

#### Critical social learning without reproductive trade-offs

We can apply our framework to study the evolution of critical social learning as considered previously (Enquist et al., 2007; Rendell et al., 2010). Assuming that learning has no marginal cost (i.e., *γ*_1_ = 1), that there is no vertical transmission (*v* = 0), and that all individuals engage into learning (*θ*_L_ = 1), we obtain that the singular IL strategy, which is given by eq. 36, reduces to

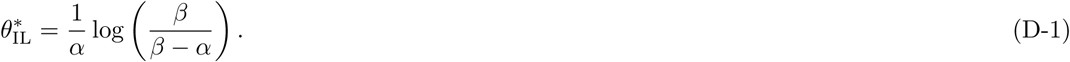

This in turn implies the level of adaptive information, which is given by eq. 39, is

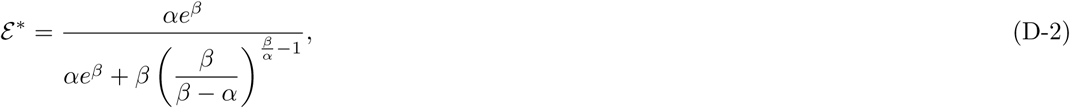

which tends to one as *β* → ∞. Using eq. (C-6), it is possible to express equilibrium eq. (D-2) in terms of 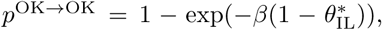 the probability of obtaining the correct phenotype via SL from an individual with the correct phenotype, and 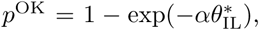 the probability of obtaining the correct phenotype via individual learning,

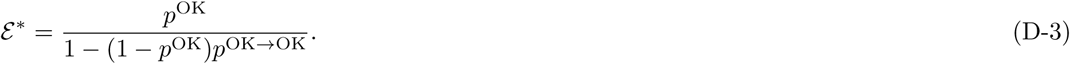

This shows immediately that *ɛ\** goes from *p*^OK^ to 1 as *p*^OK→OK^ goes from zero to one, as found by Enquist et al. (2007).

#### Cumulative culture with reproductive trade-offs

We can also use our framework to recover the results about vertical transmission obtained by Kobayashi et al. (2015). As implied by eqs. (1)–(2) of Kobayashi et al., 2015, the contributions to culture from IL and SL are now given by

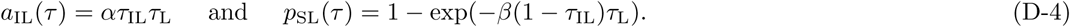

Then, to obtain a haploid version of cultural dynamics from our model, we simply drop the factor of diploid reproduction in eq. (24) and write the dynamics of cultural information as

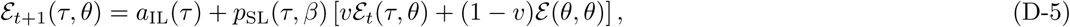

which at the equilibrium is

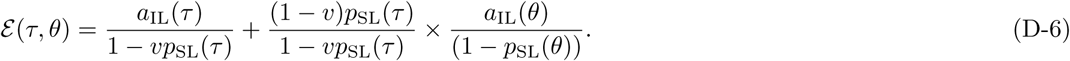

According to eq. (4) of Kobayashi et al., 2015, fecundity is given by

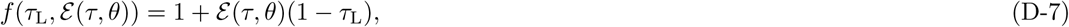

which substituted into fitness (eq. 20) with eqs. (D-4) and (D-6), give that singular strategies are

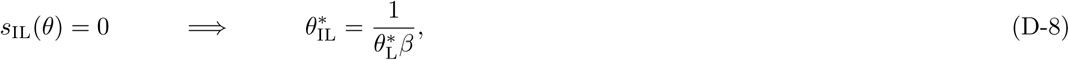

and

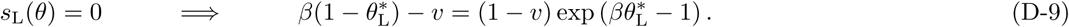

On substituting eq. (D-8) into the monomorphic equilibrium *ε*(*θ, θ*) obtained from eq. (D-6) we further have

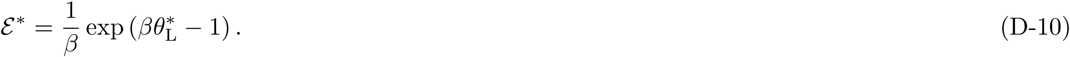

The latter three equations are equivalent to eqs. (7a)-(7c) of Kobayashi et al. (2015), respectively.

The strong effect of pure vertical transmission observed in Kobayashi et al. (2015)’s study can be seen by considering the amplification factor (Λ(*θ*), eq. 31) for learning benefits on the learning selection gradient (see eq. 29) for their model (using eq. D-5 into eq. 17),

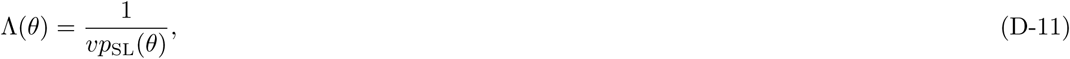

which tends to infinity as *v* and *p*_SL_(*θ*) go to one. By contrast, in our model, this amplification cannot exceed 2. Note that benefits could also be infinitely amplified in diploids if vertical transmission only occurred between mutants in mutant families so that eq. (D-5) holds.

### Appendix E: individual-based simulations

We used Mathematica (Wolfram Research, 2016) to carry out individual-based simulation of the joint evolution of IL and SL in a population with *N* individuals for our explicit model, see section 2.5 (Mfile available on request). Each individual in the population is characterized by an amount of adaptive information and two linked diploid loci that respectively determine additively the level of SL and Il performed by the individual. At the beginning of a generation (taken as stage (1) of the life cycle, see section 1.1), we calculate the fecundity *f* of each individual according to trait values and cultural variable (eq. 35). Since we assumed semelparous reproduction (eq. 20) with constant population size, we then apply a Wright-Fisher reproductive process for diploids (Ewens, 2004). Namely, we form *N* mating pairs by sampling with replacement 2 *N* individuals from the parental generation proportionally to their fecundity (hence the number of offspring produced by individual with fecundity *f* follows a binomial distribution with parameters *N* and 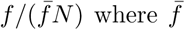 where 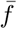 is the mean fecundity in the population). Each mating pair produces an offspring that inherits a haplotype from each parent (randomly sampled in each parent, i.e., we assume random segregation). Each locus mutates and a mutation has an additive effect sampled from a Normal distribution with mean 0 and standard deviation *σ*. Quantitative loci values are truncated to remain between 0 and 1. Then we calculate the adaptive information of each offspring according to eq. (24). At the start of a simulation, the population is initially monomorphic for SL, IL and adaptive information value.

In order to produce Fig. 4, we started with a population set for a value of 0.5 at each loci and adaptive information. We set high mutation effects with *σ* = 0.01 in order to speed up convergence. After a burn-in period of 5000 generations, we recorded the population average phenotypic values at each generation for a further 5000 generations. The temporal means and standard deviations are displayed in Fig. 4.

To test for evolutionary branching (Fig. 6 and Supplementary Figure 1), we started with a population at the evolutionary convergence stable singular values for *θ*_L_ and *θ*_IL_ and corresponding equilibrium adaptive information (eq. 26). We set low mutation effects *σ* = 0.001 and large population size *N* = 10000 as low population size prevents evolutionary branching (e.g., Wakano and Iwasa, 2013; Dèbarre and Otto, 2016). The population evolved for 1.5 *×* 10^5^ generations.

### Appendix F: numerical analysis when *λ* > 0

We here detail the numerical analyzes we performed for the case where *λ* > 0 (section 2.5.2). First, we numerically studied which points are convergence stable by holding = 2 and varying the other parameters by considering all combination of *v* = 0, 0.25, 0.5, 0.75., 1, *β* = 2, 3, 5, 9, 12, *γ*_1_ = 0, 0.1, 0.3, 0.7, 0.9, *γ*_2_ = 1, 1.5, 2, 3, 5, and *λ* = 0.1, 0.3, 0.5, 0.7, 1 (a total of 3125 parameter combinations). To do this, we iterated

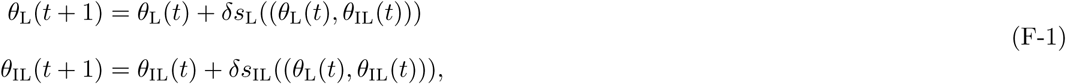

using eqs. (29) and (32) with eqs. (34)–(35) and *δ* = 0.02 until the euclidean distance between two iterates was less than 10^−5^, i.e.,

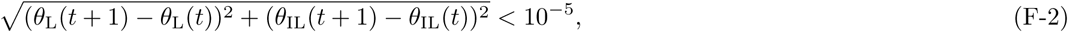

or for a maximum of 10^5^ steps. We started with 9 different starting values (*θ*_L_(0), *θ*_IL_(0)), with *θ*_L_(0) = 0.05, 0.5, 0.95 and *θ*_IL_(0) = 0.05, 0.5, 0.95. For each final values, we computed the eigenvalues of the Hessian matrix to assess local uninvadibility (Appendix B).

We find four outcomes for the convergence stability of *θ*_L_ and *θ*_IL_. (1) *θ*_L_ always converges to zero, in which case selection on *θ*_IL_ vanishes (1134/3125 cases). (2) There is a bistability and *θ*_L_ converges to either zero or to an interior value 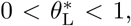 while individual learning *θ*_IL_ goes to one (i.e., no social learning, 689/3125 cases). (3) There is a bistability and *θ*_L_ converges to either zero or both *θ*_L_ and *θ*_IL_ converge to an interior equilibrium 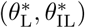 where *θ*_IL_ satisfies eq. (36) (1296/3125 cases, see Fig 5). (4) In very few cases (6/3125 cases), there are three convergence stable equilibria: either *θ*_L_ converges to zero; or *θ*_L_ converges to an interior value 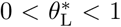 while individual learning *θ*_IL_ goes to one; or both *θ*_L_ and *θ*_IL_ converge to an interior equilibrium 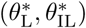 (where *θ*_IL_ satisfies eq. (36)).

When we assess the local uninvadability (using the eigenvalues of the Hessian matrix, Appendix B) of interior convergence stable equilibria 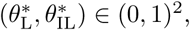 we find that they are also locally uninvadable in the majority of cases (1290 out of 1302). In 12 cases (out of 1302), the interior convergence stable equilibria 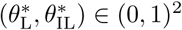 was locally invadable, suggesting that learning undergoes disruptive selection at these equilibria. By looking at the parameter values under which disruptive selection occurs, we find that greater rates of vertical transmission (*v*) disfavor disruptive selection. For instance, when *α* = 2, *β* = 12, *γ*_1_ = 0.7, *γ*_2_ = 5, *λ* = 0.7, the convergence stable point 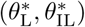 is only uninvadable when *v >* 0.4 (Supplementary Figure 1). Individual based simulations confirmed our analysis of disruptive selection, and showed that disruptive selection leads to evolutionary branching and the emergence of a polymorphism that can be maintained in the long run in some cases (Supplementary Figure 1).

---

**Supplementary Figure 1:**
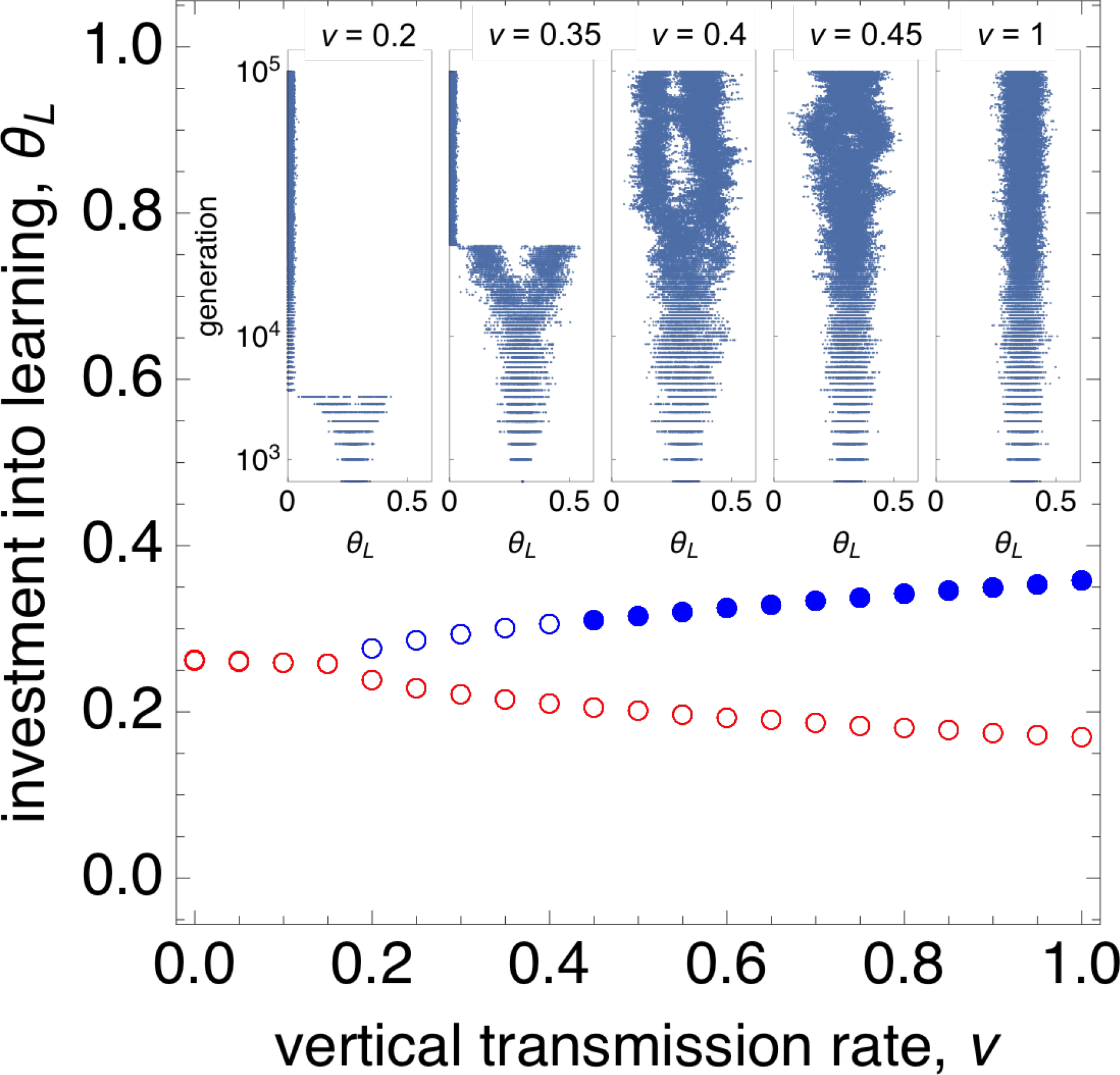
Evolutionary branching and vertical transmission. Outset: Interior singular learning strategies of *θ*_L_ according to vertical transmission rate, generated by computing the roots of the selection gradients (eqs. 29 and 32, other parameters: *α* = 2, *β* = 12, *γ*_1_ = 0.7, *γ*_2_ = 5, *λ* = 0.7). For each singular value, we computed their convergence stability and local invadability (Appendix B). Red empty circles indicate singular strategies that are convergence unstable and thus represent threshold values above which learning is favored by selection. Blue empty circles are convergence stable but invadable (i.e., potential evolutionary branching points), while blue filled circles are convergence stable and uninvadable (i.e., ESSs). Hence, the analytical model predict that for 0.4 ≤ *v* ≤ 1 selection is stabilizing, for 0.2 ≤ *v* ≤ 0.4 selection is disruptive, while for *v <* 0.2 there is no interior convergence stable point. Inset: values of *θ*_L_ for each individual in the population for every 500 generations under individual-based simulations started at the convergence stable singular value, where different panels correspond to different *v* values (Appendix E). As predicted by the model, we see that when *v >* 0.4, the population remains unimodal, when 0.2 ≤ *v* ≤ 0.4, disruptive selection occurs, which results in evolutionary branching whereby the population splits into two morphs. However, when *v* decreases too much below 0.4, the two morphs are not maintained in the long run and the population converges to zero learning. This is because evolutionary branching causes the population to cross the threshold value of learning, resulting in the collapse of learning.

